# Reciprocal Control of Motility and Biofilm Formation by the PdhS2 Two-Component Sensor Kinase of *Agrobacterium tumefaciens*

**DOI:** 10.1101/148429

**Authors:** Jason E. Heindl, Daniel Crosby, Sukhdev Brar, Tiyan Singletary, Daniel Merenich, Justin L. Eagan, Aaron M. Buechlein, Eric L. Bruger, Christopher M. Waters, Clay Fuqua

## Abstract

A core regulatory pathway that directs developmental transitions and cellular asymmetries in *Agrobacterium tumefaciens* involves two overlapping, integrated phosphorelays. One of these phosphorelays putatively includes four histidine sensor kinase homologues, DivJ, PleC, PdhS1, and PdhS2, and two response regulators, DivK and PleD. In several different alphaproteobacteria, this pathway influences a conserved downstream phosphorelay that ultimately controls the phosphorylation state of the CtrA master response regulator. The PdhS2 sensor kinase reciprocally regulates biofilm formation and swimming motility. In the current study the mechanisms by which the *A. tumefaciens* sensor kinase PdhS2 directs this regulation are delineated. PdhS2 lacking a key residue for phosphatase activity is markedly deficient in proper control of attachment and motility phenotypes, whereas a kinase-deficient PdhS2 mutant is only modestly affected. A genetic interaction between DivK and PdhS2 is revealed, unmasking one of several connections between PdhS2-dependent phenotypes and transcriptional control by CtrA. Epistasis experiments suggest that PdhS2 can function independently of the CckA sensor kinase, the cognate sensor kinase for CtrA which is inhibited by DivK. PdhS2 dynamically localizes to the daughter cell pole in dividing cells. Global expression analysis of the *pdhS2* mutant reveals a restricted regulon, functioning through CtrA to separately control motility and regulate levels of the intracellular signal cyclic diguanylate monophosphate (cdGMP), thereby affecting production of adhesive polysaccharides and attachment. We hypothesize that in *A. tumefaciens* the CtrA regulatory circuit has expanded to include additional inputs through addition of PdhS-type sensor kinases, likely fine-tuning the response of this organism to the soil microenvironment.

**IMPORTANCE:** Bacterial developmental processes, such as morphological transformations and behavioral transitions, are tightly regulated. In many alphaproteobacteria cell division and development are coordinated by a specific suite of conserved histidine kinases and their partnered regulatory proteins. Here we describe how the histidine kinase PdhS2 of *Agrobacterium tumefaciens* regulates complex phenotypes including biofilm formation and motility. PdhS2 genetically interacts with a single-domain response regulator, DivK, and the intracellular signal cyclic diguanylate monophosphate. PdhS2 dynamically localizes to the new pole of recently divided cells, contributing to the regulatory processes that dictate whether these cells remain motile or initiate biofilm formation. These findings expand our understanding of the complex network that integrates cell division and developmental control in *A. tumefaciens* and related alphaproteobacteria.

## INTRODUCTION

Bacteria are sometimes considered to be elementary life forms, with simple body plans, streamlined reproductive cycles, and monolithic behavior when compared with higher eukaryotes. To the contrary, many bacteria can exhibit a remarkable diversity of developmental complexity, both temporal and morphological (1, 2). Even bacterial species whose cells appear morphologically uniform, such as rod-shaped *Escherichia coli* or coccoid *Staphylococcus aureus,* possess distinct cellular architectures as well as intricately timed cell division programs, and a large number of bacteria can form multicellular biofilms (3, 4). Developmental processes in bacteria, as in higher eukaryotes, are driven by factors that may be considered both cell-intrinsic and cell-extrinsic. Intrinsic factors include genomic and proteomic content, while extrinsic factors comprise environmental conditions, such as pH and temperature, which cells sense and to which they respond (5).

Members of the *Alphaproteobacteria* group include host-associated pathogens (*e.g. Brucella* sp., *Bartonella* sp.), host-associated commensals (*e.g. Sinorhizobium* sp., *Bradyrhizobium* sp.), and free-living aquatic and marine bacteria (*e.g. Caulobacter* sp., *Rhodobacter* sp., *Ruegeria* sp.). It is now recognized that several alphaproteobacteria divide asymmetrically, during which cells elongate, duplicate and segregate their genomic content between two non-equivalent compartments of predivisional cells, and finally generate two cells by cytokinesis (6, 7). Notably, cellular components are unevenly distributed between the two daughter cells during cell division, including surface structures (*e.g.* flagella and polar polysaccharides), cell wall components (*e.g.* peptidoglycan), and even cytoplasmic complexes (*e.g.* heat shock proteins). For example, there may be a clear segregation of existing organelles to one daughter cell while the second cell generates these structures *de novo* (6, 8–10). Although the specific details may vary among different taxa, the end result is the production of a young daughter cell and a comparatively aging mother cell. Not only does this uneven division partition senescence among the products of cell division, but it also allows for the generation of functionally distinct cell types. For example, in *Caulobacter crescentus* the non-motile stalked cell type can attach to surfaces using its polar adhesin called the holdfast (11). This stalked cell then serves as the mother cell during multiple rounds of cell division, generating and releasing motile swarmer cells upon each cytokinetic event (12). Motile swarmer cells are prohibited from entering the cell division cycle until differentiation into the non-motile stalked form (13, 14).

Underlying asymmetric cell division is subcellular differentiation that includes localization of specific regulatory proteins to programmed locations within each cell (15). Prominent among these in many alphaproteobacteria are components of two overlapping phosphorelays, the first which functions through the response regulators DivK and PleD (the DivK-PleD relay) and the second which functions primarily through the response regulator CtrA (the CtrA relay). The pathways are connected through DivK, which controls initiation of the CtrA relay by regulating its cognate sensor kinase CckA (16, 17). Collectively we refer to these two relays as the DivK-CtrA pathway. In the well-studied *C. crescentus* system the membrane-associated sensor histidine kinases PleC and DivJ control the phosphorylation state of DivK and PleD, and localize to opposing poles of the predivisional cell (18–21). Through antagonistic kinase and phosphatase activities on DivK and PleD, their target response regulators, PleC and DivJ inversely manifest their activity on the most downstream component of the DivK-CtrA pathway, the response regulator CtrA (22–25). DivJ is retained at the stalked cell pole and serves as a DivK/PleD kinase, increasing the DivK~P concentration and diminishing CtrA~P levels in this region of the cell (Fig. 1A). Conversely, PleC localizes to the pole distal to the stalk, where the single polar flagellum is assembled, dephosphorylating DivK, leading to increased CtrA~P levels and activity. Phospho-CtrA binds to the replication origin thereby preventing DNA replication, and also acts as a transcriptional regulator for many genes, including activating those for assembly of the flagellum and motility (26–28). The CtrA relay is also influenced by the DivK-PleD relay through levels of the second messenger cyclic diguanylate monophosphate (cdGMP). DivJ-dependent phosphorylation of PleD at the stalk pole of the predivisional cell stimulates its diguanylate cyclase activity, resulting in higher levels of cdGMP at this end of the cell. The CckA kinase that initiates the CtrA relay is also biased away from its kinase and towards its phosphatase activity by direct allosteric control through high levels of cdGMP, thereby reinforcing a CtrA~P gradient, relatively low at the stalk pole and increasing towards the distal pole (29–34) (Fig. 1A).

**Figure 1. The PdhS kinases of *C. crescentus* and *A. tumefaciens* differentially localize and affect phenotypic outputs through response regulators DivK, PleD, and CtrA.** (A) Cartoon model of known localization of the namesake PdhS kinases from *C. crescentus*, PleC and DivJ, and three PdhS kinases from *A. tumefaciens*, PdhS1, PdhS2, and DivJ. Kinases represented as colored ovals with black border experimentally localize to the indicated poles. The PleC oval without a border has not been experimentally demonstrated to localize in *A. tumefaciens*. As a result of this localization phosphorylation status of direct PdhS kinase targets, DivK and PleD, and the indirect target, CtrA, may be differentially affected. (B) Multiple sequence alignment of the HisKA domain from the PdhS kinases of *C. crescentus* and *A. tumefaciens*. Sequences were aligned using the Clustal Omega web service hosted by the European Molecular Biology Laboratory (EMBL)-European Bioinformatics Institute. The four PdhS kinases from *A. tumefaciens* plus PleC and DivJ from *C. crescentus* were included. Also included are two additional predicted PdhS kinases, CC_0652 and CC_1062, from *C. crescentus*. The EnvZ sensor kinase is included for comparison. Yellow highlighting indicates residues that define the PdhS kinases. The conserved histidine and threonine residues mutated in this work are in bold.

*Agrobacterium tumefaciens* is a plant pathogen of the *Alphaproteobacteria* class that is not stalked, but like *C. crescentus*, divides asymmetrically generating a motile daughter cell from a mother cell (6). As a facultative pathogen, the *A. tumefaciens* lifestyle substantially differs from that of the freshwater oligotroph *C. crescentus*. Nonetheless core components of the DivK-CtrA pathway are well conserved in *A. tumefaciens*, including three non-essential <u>P</u>leC/<u>D</u>ivJ <u>h</u>omologue <u>s</u>ensor kinase (PdhS) homologues PleC (Atu0982), PdhS1 (Atu0614), and PdhS2 (Atu1888). The *divJ* gene (Atu0921) is essential in *A. tumefaciens* (35, 36). We have previously shown that the three non-essential PdhS homologues have distinct roles in the normal cellular development of *A. tumefaciens* (35). Mutants in PleC and PdhS1, as well as the *A. tumefaciens* DivK homologue, all manifested marked effects on cell division, with branched and elongated cells, as well as deficiencies in motility and biofilm formation. To date the essentiality of *divJ* has precluded exhaustive phenotypic analysis of its role (35). The fourth PdhS family member, PdhS2, does not appear to participate in regulation of cell division as all cells are morphologically wild-type in appearance (35). Loss of PdhS2, however, results in dramatically increased attachment and biofilm formation, and a simultaneous dramatic reduction in motility. Reciprocal regulation of these phenotypes is often a hallmark of regulation by cdGMP (37). In this work we further explore the mechanism by which PdhS2 regulates attachment and motility. Our results genetically connect DivK with PdhS2 and transcriptional profiling clearly implicates CtrA as their downstream regulatory effector. We also show a clear intersection of PdhS2 activity and the activity of several diguanylate cyclases, suggesting that PdhS2 and cdGMP coordinately regulate biofilm formation and motility in *A. tumefaciens*. Finally, we observe that PdhS2 dynamically localizes to the newly generated poles of both the new daughter cell and the mother cell following cytokinesis. Collectively, our findings suggest that PdhS2 activity is specifically required for proper development of motile daughter cells.

## RESULTS

### Mutational analysis of PdhS2 reveals coordinate regulation through kinase and phosphatase activities

Members of the PdhS family of sensor histidine kinases contain a conserved HATPase_c catalytic domain at their carboxyl termini and an upstream conserved HisKA dimerization/phosphoacceptor domain. Many sensor kinases exhibit bifunctional catalytic activity, alternately acting as kinase or phosphatase, and *C. crescentus* PleC is one such example (18, 22, 38). Multiple sequence alignment of the HisKA domain from the *A. tumefaciens* and *C. crescentus* PdhS family kinase homologues reveals a high level of conservation of this domain including the phospho-accepting histidine residue (H271 of PdhS2) and a threonine residue predicted to be important for phosphatase activity (T275 of PdhS2) (Fig. 1B).

To test the requirement of the conserved phospho-accepting histidine for PdhS2 activity we mutated this residue to alanine (H271A). Ectopic expression of PdhS2^H271A^ (plasmid-borne *P*_lac_-*pdhS2*, kinase-negative, K; phosphatase-positive, P^+^) effectively complemented the attachment and motility phenotypes of the Δ*pdhS2* mutant similar to wild type *pdhS2* (Fig. 2A). These data indicate that this histidine residue is not crucial for PdhS2 regulation of swimming motility and biofilm formation. When *pdhS2* is ectopically expressed in the wild type, it causes a slight but significant stimulation of biofilm formation, and the *pdhS2*^H271A^ (K^−^P^+^) mutation reverses this effect.

**Figure 2. Evaluation of roles for PdhS2 kinase and phosphatase activities and genetic interactions with *divK*.** (A) The ability of plasmid-borne wild-type PdhS2 (p-*pdhS2*), the kinase-null allele (p-*pdhS2*, K^−^P^+^), or the phosphatase-null allele (p-*pdhS2,* K^+^P^−^) to complement the Δ*pdhS2* biofilm formation (black bars) and swimming motility (white bars) phenotypes was evaluated using *P*_*lac*_-driven expression of each allele. Static biofilm formation was measured after 48 h (black bars) and swim ring diameter after 7 days (white bars). Adherent biomass on PVC coverslips was determined by adsorption of crystal violet. Crystal violet was then solubilized and A_600 nm_ values were normalized to culture density (OD_600_). Data are the mean of three independent experiments each of which contained three technical replicates (N = 3). Swim ring diameters were measured after single-colony inoculation into low density swim agar and incubation at room temperature. Data are the mean of nine independent experiments (N = 9). (B) Biofilm formation (black bars) and swimming motility (white bars) were evaluated in the indicated strains. Experiments were performed and data analyzed as described for (A) above. (C) The effect of plasmid-borne wild-type PdhS2 (p-*pdhS2*), the kinase-null allele (p-*pdhS2* (K^−^P^+^)), or the phosphatase-null allele (p-*pdhS2* (K^+^P^−^)) on biofilm formation (black bars) and swimming motility (white bars) when expressed from the *P*_lac_ promoter in the Δ*divK* mutant background was evaluated as in (A) and (B) above. For presentation all data are normalized to WT and expressed as %WT ± standard error of the mean (S.E.). (^a^) = *P* < 0.05 compared to wild-type strain or wild-type strain carrying empty vector. (^b^) = *P* < 0.05 compared to Δ*pdhS2* strain carrying empty vector (A), or compared to the Δ*pdhS2* mutant strain (B), or compared to the Δ*divK* strain carrying empty vector (C). Statistical significance was determined using Student’s *t* test.

Efficient phosphatase activity of many sensor kinases requires a conserved threonine residue roughly one α-helical turn (4 residues) downstream of the phospho-accepting histidine residue (39, 40). We therefore mutated this conserved threonine residue to alanine (Thr275A). In contrast to the PdhS2^H271A^ mutant protein (K^−^P^+^), equivalent ectopic expression of the PdhS2^T275A^ allele (K^+^P^−^) failed to complement the Δ*pdhS2* motility and attachment phenotypes, and in fact exacerbated them (Fig. 2A). When expressed in wild type, the PdhS2^T275A^ allele (K^+^P^−^) caused modest stimulation of biofilm formation and slightly decreased motility. A double mutant allele of PdhS2 with both the histidine and threonine residues mutated (K^-^P^-^) had no effect on these phenotypes (Fig. S1). Together these results suggest that it is the balance of kinase and phosphatase activity that dictates PdhS2 control over its targets, with the kinase stimulating biofilm formation and decreasing motility, and the phosphatase activity diminishing biofilm formation and promoting motility. The phosphatase activity however, appears to play the dominant role under laboratory culture conditions.

### Mutations in *divK* are epistatic to *pdhS2* mutations

Members of the PdhS family of sensor kinases were originally identified based on homology with their namesakes DivJ and PleC of *C. crescentus* (41, 42) (Fig. 1B). Based on this homology all PdhS family members are predicted to interact with the single domain response regulator DivK (42) and also interact with the diguanylate cyclase response regulator PleD. Prior work from our laboratory has shown that both swimming motility and adherent biomass are diminished in the Δ*divK* mutant, implying that DivK activity is required for proper regulation of these phenotypes in *A. tumefaciens* (35). In contrast, PdhS2 inversely regulates these phenotypes; a Δ*pdhS2* mutant is non-motile but hyperadherent. To determine whether PdhS2 genetically interacts with DivK we constructed a Δ*divK*Δ*pdhS2* mutant and compared swimming motility and biofilm formation in this strain to wild-type and parental single deletion strains (Fig. 2B). As reported, loss of either *divK* or *pdhS2* reduced swimming motility as measured by swim ring diameter on motility agar. Biofilm formation on PVC coverslips in the Δ*divK* mutant was diminished relative to the wild-type C58 strain while for the Δ*pdhS2* mutant it was dramatically increased. The Δ*divK*Δ*pdhS2* mutant was similar to the Δ*divK* mutant in both assays, with no significant difference in the efficiency of either swimming motility or biofilm formation between the two strains. These data support the proposed genetic interaction between *divK* and *pdhS2*, with the *divK* mutation epistatic to *pdhS2* for biofilm formation and swimming motility.

Swim ring diameters of the Δ*divK* and Δ*divK*Δ*pdhS2* mutants were decreased by ~20% compared to wildtype whereas the decrease in Δ*pdhS2* swim ring diameters was ~40% compared to wildtype, suggesting that the nature of the defect in swimming motility differs between these two classes of mutants and that loss of *divK* partially restores motility in the absence of *pdhS2*. Indeed, it was earlier noted that although both the Δ*divK* and the Δ*pdhS2* single deletion mutants produce polar flagella, very few Δ*pdhS2* mutant bacteria were observed to be motile under wet-mount microscopy implying that the swimming defect is due to diminished flagellar activity rather than flagellar assembly (35). The Δ*divK* mutant, however, was readily observed to be motile under wet-mount microscopy. Similarly, the Δ*divK*Δ*pdhS2* mutant generates polar flagella and its motility is readily observed under wet-mount microscopy. Both the Δ*divK* and Δ*divK*Δ*pdhS2* mutants, and not the Δ*pdhS2* mutant, generate aberrant cell morphologies including elongated and branched cells (35) (Fig. S2).

Further support for a genetic interaction between PdhS2 and DivK was provided by plasmid-borne, wild-type PdhS2 expressed ectopically from a *P*_lac_ promoter. Induced expression of *pdhS2* rescues swimming motility and returns biofilm formation closer to wild type levels in the Δ*pdhS2* mutant, albeit incompletely (Fig. 2A). However, as predicted from the Δ*divK*Δ*pdhS2* phenotypes, plasmid-borne provision of PdhS2 in the Δ*divK* mutant had no significant effect on either biofilm formation or swimming motility (Fig. 2C). Expression of either the kinase-null or the phosphatase-null allele of PdhS2 in the Δ*divK* background similarly had no effect on biofilm formation or swimming motility (Fig. 2C).

### Kinase-locked allele of CckA does not suppress *pdhS2* phenotypes

One key PdhS target among multiple bacterial taxa is the hybrid histidine kinase CckA. CckA exhibits dynamic regulation dependent upon both phosphorylation status of DivK and local levels of cdGMP (31). CckA ultimately serves as either a source or a sink for CtrA phosphorylation through a phosphorelay that includes the histidine phosphotransferase ChpT (29). Previously we identified a mutation in CckA that results in a kinase-locked allele which is insensitive to regulation by DivK and cdGMP, CckA^Y674D^ (31, 35). Expression of this allele in the Δ*pleC* background suppressed the swimming motility defect of the Δ*pleC* strain but had no effect on swimming motility in the Δ*divK* background (35). These results were consistent with PleC activity proceeding through DivK and with DivK negatively regulating CckA kinase activity. We reasoned that if PdhS2 similarly functioned through DivK and CckA, expression of CckA^Y674D^ in the Δ*pdhS2* background would suppress the motility and biofilm phenotypes of this strain. However, induced expression of wild-type CckA or the CckA^Y674D^ allele only marginally impacted these phenotypes (Fig. S3 and S4). These observations suggest that PdhS2 functions differently than PleC.

### PdhS2 and DivJ localize to the pole of *A. tumefaciens*

A mechanism for establishing and maintaining developmental asymmetries in bacteria is the differential polar localization of proteins with opposing functionalities (43, 44). Several members of the PdhS family of sensor kinases localize to one or both bacterial poles (18, 22, 45). Using a full-length PdhS2-GFP fusion that retains wild type functionality, expressed from *P*_*lac*_ on a low copy number plasmid, we tracked localization of PdhS2 in *A. tumefaciens* following IPTG induction (Fig. 3A). In both wild-type (data not shown) and Δ*pdhS2* backgrounds PdhS2-GFP localized primarily to the new pole of predivisional cells. Time-lapse microscopy of the Δ*pdhS2* mutant expressing PdhS2-GFP revealed apparent dynamic relocalization of PdhS2-GFP to the newly generated pole of the daughter cell coincident with cytokinesis. In the mother cell a new focus of PdhS2-GFP subsequently accumulates at the new pole generated following septation and daughter cell release. These time-lapse experiments clearly indicate that PdhS2-GFP localizes to the actively growing pole of the cell, and is lost at that pole as it developmentally matures and the cell proceeds to the predivisional state. Following cytokinesis PdhS2-GFP localizes to the newly generated, younger poles of both the daughter cell and the mother cell. This dynamic localization is consistent with PdhS2 activity being restricted to the motile-cell compartment of predivisional cells and to newly generated motile cells.

**Figure 3. PdhS2 and DivJ are polarly localized in *A. tumefaciens*.** Time-lapse microscopy of a C-terminal green fluorescent protein fusions to PdhS2 (A) and DivJ (B). Overlaid phase and fluorescent images, acquired sequentially on Nikon E800 fluorescence microscope with a CCD camera using the 100 × objective. Time between panels is 40 minutes. To the right of each image is a cartoon interpretation of the image.

Since PdhS2 localizes to the new pole of dividing cells and primarily requires its phosphatase activity there likely exist one or more old pole-localized kinases opposing PdhS2 activity. The most obvious candidate for this is DivJ, which localizes to the old pole in *C. crescentus* and acts as a DivK kinase in both *C. crescentus* and *S. meliloti* (42, 46). Time-lapse microscopy of a full length DivJ-GFP fusion in wild-type *A. tumefaciens* reveals localization to the old pole in mother cells that is not redistributed over the course of multiple cell division cycles (Fig. 3B). Our findings are confirmed by a recent report of similar localization patterns for PdhS2 and DivJ, which also demonstrates PdhS1 localization to the old pole of the bacterium, similar to DivJ (47).

### Expression of predicted CtrA-dependent promoters

The known architecture of the DivK-CtrA pathway predicts that PdhS2 impacts developmental phenotypes through the transcriptional regulator CtrA. In *C. crescentus*, CtrA is known to directly regulate at least 55 operons, acting as either an activator or repressor of transcription, and to control DNA replication (22, 28). *A. tumefaciens* CtrA is predicted to act similarly, binding to DNA in a phosphorylation-dependent manner and regulating DNA replication and transcription. *A. tumefaciens* CtrA is 84% identical to *C. crescentus* CtrA at the amino acid level and purified *C. crescentus* CtrA binds to a site upstream of the *A. tumefaciens ccrM* gene (48). Furthermore, computational analysis of multiple alphaproteobacterial genomes uncovered numerous cell cycle regulated genes preceded by a consensus CtrA binding site (49). We therefore evaluated CtrA activity by examining the transcription of several known and predicted CtrA-dependent promoters from both *C. crescentus* and *A. tumefaciens* in wild-type, Δ*pdhS2*, and Δ*divK A. tumefaciens* strain backgrounds. The *ccrM*, *ctrA*, and *pilA* promoters from *C. crescentus* were chosen to represent CtrA-activated promoters likely to be similarly regulated in *A. tumefaciens* (13, 14, 50, 51). In the Δ*pdhS2* background, expression levels from both the *ctrA* and *pilA* promoters from *C. crescentus* were significantly reduced while transcription from the *C. crescentus ccrM* promoter was unchanged (Table 1). In the *A. tumefaciens* Δ*divK* background the *C. crescentus ccrM* and *ctrA* promoters exhibited increased activity while the *pilA* promoter was unchanged (Table 1). These data are consistent with *A. tumefaciens* CtrA regulating transcription of known CtrA-dependent promoters, and with PdhS2 and DivK inversely regulating CtrA activity in *A. tumefaciens*.

**Table 1.**
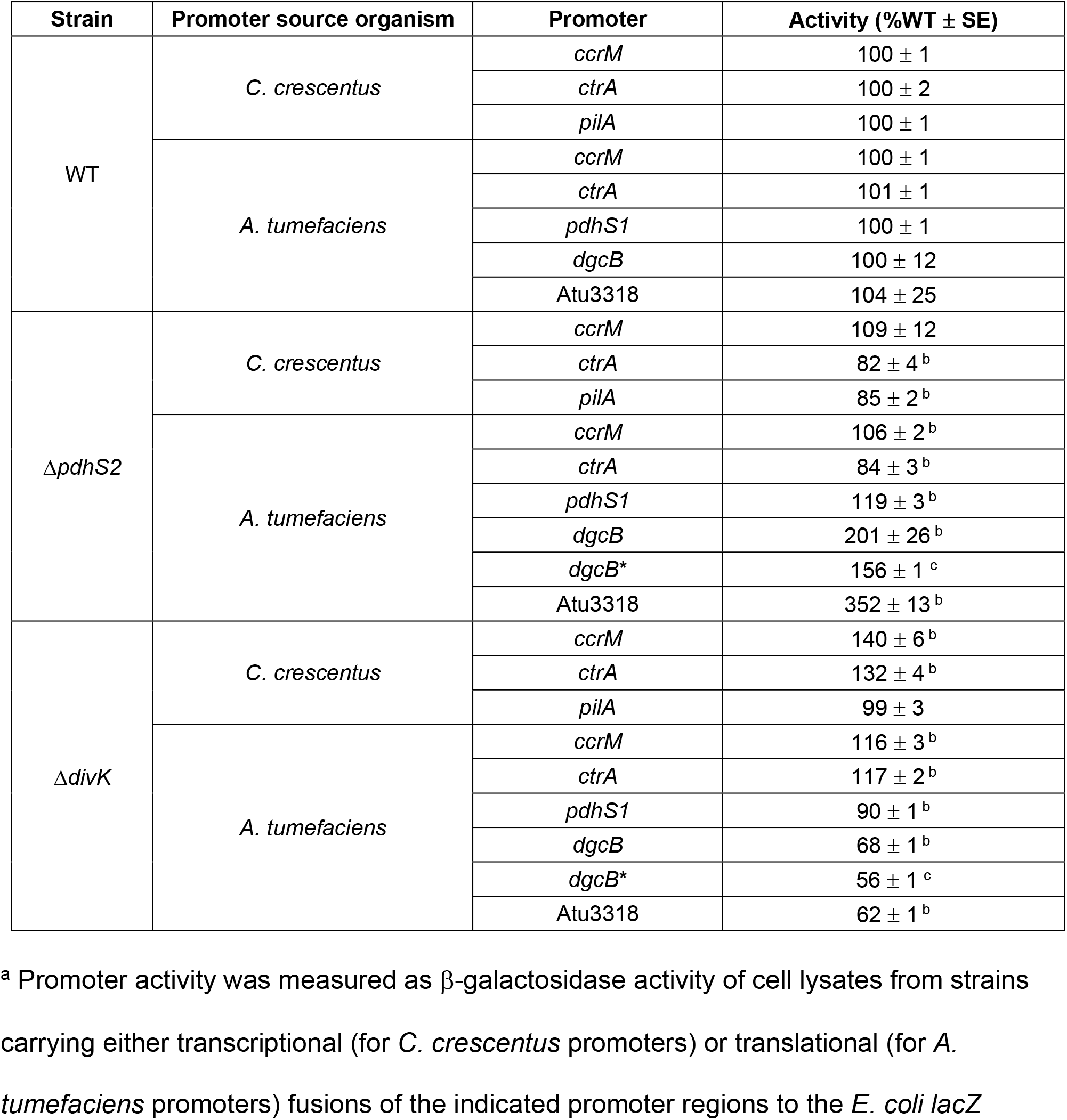
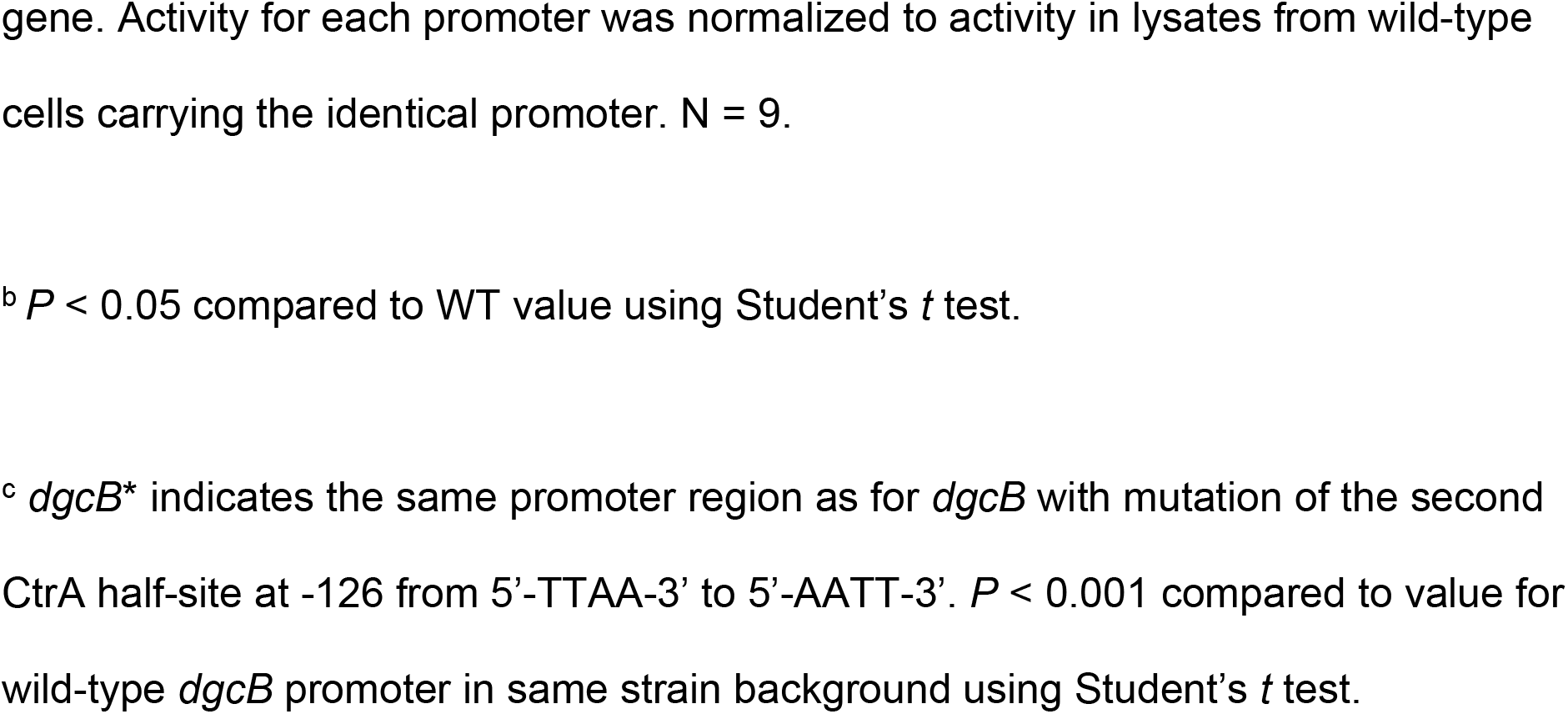
Promoter activity of selected known and predicted CtrA-dependent promoters.^a^

| Strain | Promoter source organism | Promoter | Activity (%WT ± SE) |
| --- | --- | --- | --- |
| WT | *C. crescentus* | *ccrM* | 100 ± 1 |
| | | *ctrA* | 100 ± 2 |
| | | *pilA* | 100 ± 1 |
| | *A. tumefaciens* | *ccrM* | 100 ± 1 |
| | | *ctrA* | 101 ± 1 |
| | | *pdhS1* | 100 ± 1 |
| | | *dgcB* | 100 ± 12 |
| | | Atu3318 | 104 ± 25 |
| $\Delta pdhS2$ | *C. crescentus* | *ccrM* | 109 ± 12 |
| | | *ctrA* | 82 ± 4<sup>b</sup> |
| | | *pilA* | 85 ± 2<sup>b</sup> |
| | *A. tumefaciens* | *ccrM* | 106 ± 2<sup>b</sup> |
| | | *ctrA* | 84 ± 3<sup>b</sup> |
| | | *pdhS1* | 119 ± 3<sup>b</sup> |
| | | *dgcB* | 201 ± 26<sup>b</sup> |
| | | *dgcB\** | 156 ± 1<sup>c</sup> |
| | | Atu3318 | 352 ± 13<sup>b</sup> |
| $\Delta divK$ | *C. crescentus* | *ccrM* | 140 ± 6<sup>b</sup> |
| | | *ctrA* | 132 ± 4<sup>b</sup> |
| | | *pilA* | 99 ± 3 |
| | *A. tumefaciens* | *ccrM* | 116 ± 3<sup>b</sup> |
| | | *ctrA* | 117 ± 2<sup>b</sup> |
| | | *pdhS1* | 90 ± 1<sup>b</sup> |
| | | *dgcB* | 68 ± 1<sup>b</sup> |
| | | *dgcB\** | 56 ± 1<sup>c</sup> |
| | | Atu3318 | 62 ± 1<sup>b</sup> |

<sup>a</sup> Promoter activity was measured as β-galactosidase activity of cell lysates from strains carrying either transcriptional (for *C. crescentus* promoters) or translational (for *A. tumefaciens* promoters) fusions of the indicated promoter regions to the *E. coli lacZ*
gene. Activity for each promoter was normalized to activity in lysates from wild-type cells carrying the identical promoter. N = 9.
<sup>b</sup> $P < 0.05$ compared to WT value using Student's $t$ test.
<sup>c</sup> *dgcB*<sup>\*</sup> indicates the same promoter region as for *dgcB* with mutation of the second CtrA half-site at -126 from 5'-TTAA-3' to 5'-AATT-3'. $P < 0.001$ compared to value for wild-type *dgcB* promoter in same strain background using Student's $t$ test.

From *A. tumefaciens* the *ccrM* promoter is the only promoter for which prior experimental data suggest CtrA-dependent regulation, thus this promoter was selected for analysis (48). In addition to *ccrM*, putative *A. tumefaciens* promoters for *ctrA* and *pdhS1* were selected for analysis based on the presence of at least one predicted CtrA binding site as well as hypothesized cell cycle regulation of these loci. Transcriptional activity from the *A. tumefaciens ctrA* and *pdhS1* promoter constructs showed inverse regulation in the Δ*pdhS2* and Δ*divK* backgrounds, with expression decreased from the *ctrA* promoter and increased for the *pdhS1* promoter in the Δ*pdhS2* mutant, and exactly reversed in the Δ*divK* mutant. Although absence of *pdhS2* had little effect on the *A. tumefaciens ccrM* promoter, transcription from this promoter was significantly increased in the Δ*divK* background (Table 1). These data are congruent with the above data for *C. crescentus* CtrA-dependent promoters and further support CtrA regulation of cell cycle-responsive genes in *A. tumefaciens*.

### Global transcriptional analysis of PdhS2 activity

To determine the effect of PdhS2 activity on the *A. tumefaciens* transcriptome we used whole genome microarrays. Gene expression was compared between WT and Δ*pdhS2* strains grown to exponential phase in minimal media. Of 5338 unique loci represented on the arrays 39 genes were differentially regulated above our statistical cut-offs (P values, ≤ 0.05; log_2_ ratios of ≥ ± 0.50; Table 2). Of these, 24 genes were significantly upregulated, indicating negative regulation by PdhS2. Upregulated genes included *dgcB*, previously shown to contribute to elevated biofilm formation in hyperadherent *A. tumefaciens* mutants disrupted in the motility regulators VisN and VisR (52). Also upregulated was Atu3318, encoding a LuxR-type transcription factor, that similar to *dgcB* was previously identified by its elevated levels in *visNR* null mutants. Downregulated genes in the *pdhS2* mutant included six succinoglycan biosynthetic genes, consistent with our previous results showing positive regulation of succinoglycan production by PdhS2 (35).

**Table 2.**
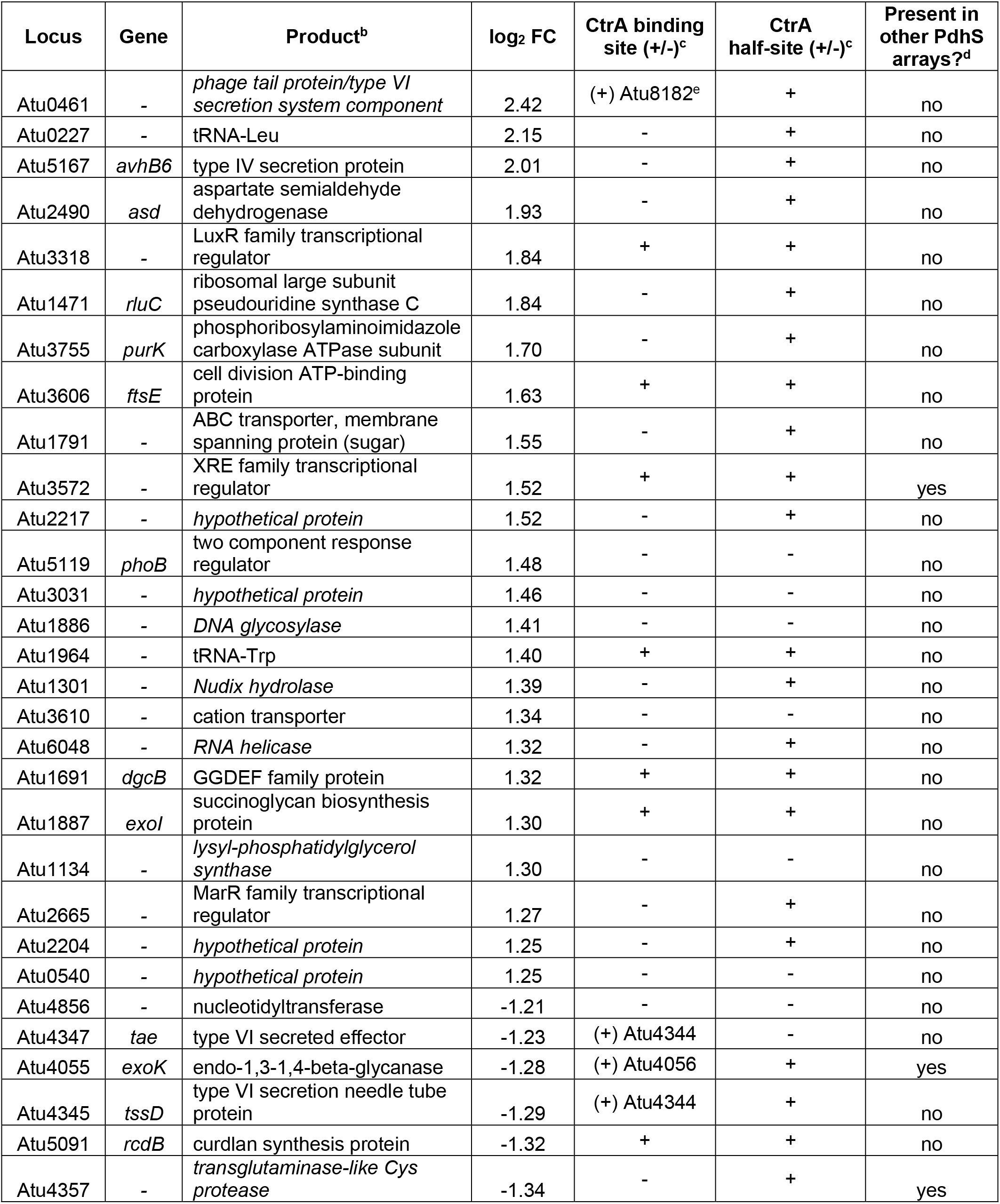
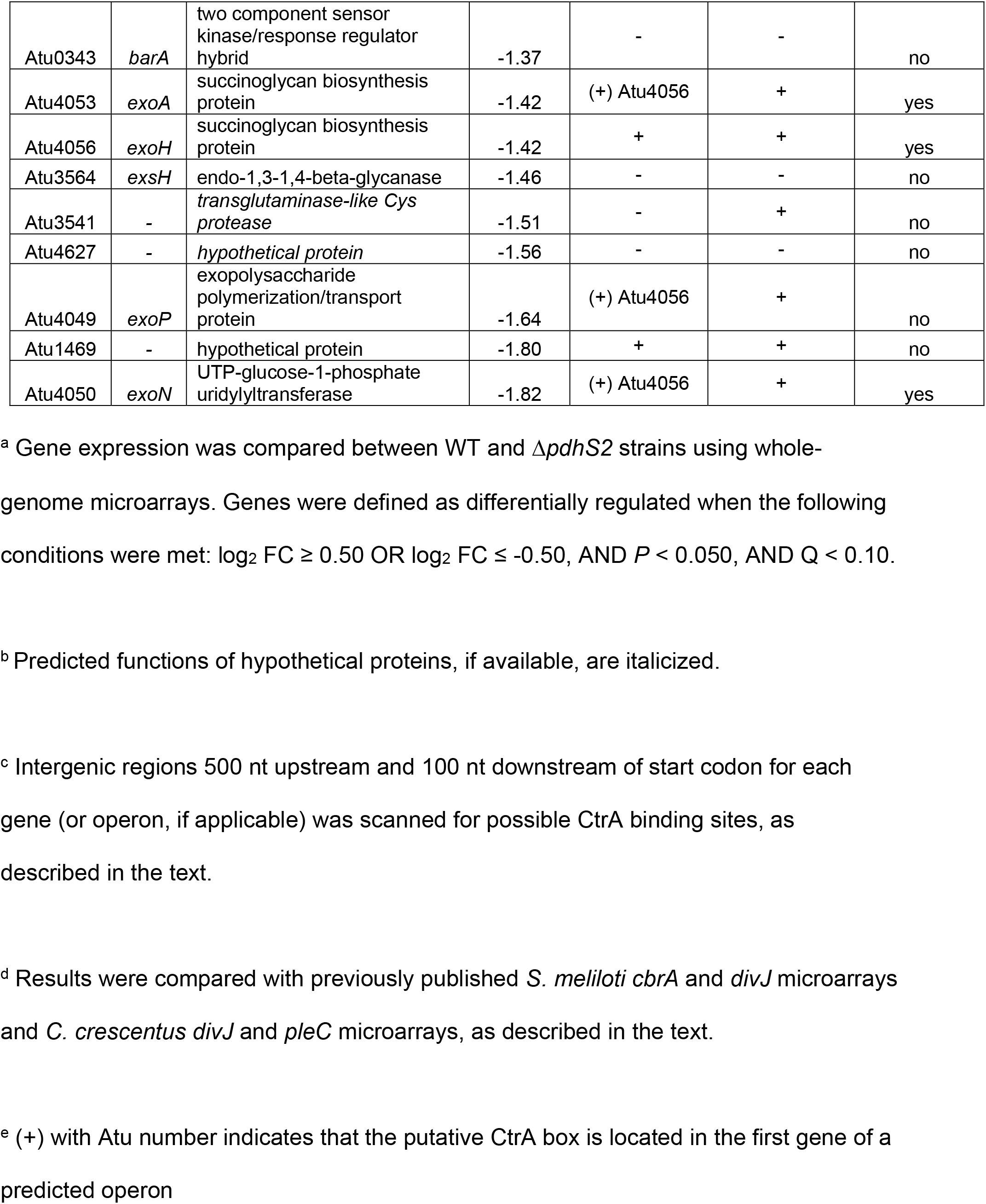
Differentially-regulated genes in the absence of *pdhS2*.^a^

| Locus | Gene | Product<sup>b</sup> | log<sub>2</sub> FC | CtrA binding site (+/-)<sup>c</sup> | CtrA half-site (+/-)<sup>c</sup> | Present in other PdhS arrays?<sup>d</sup> |
| --- | --- | --- | --- | --- | --- | --- |
| Atu0461 | - | *phage tail protein/type VI secretion system component* | 2.42 | (+) Atu8182<sup>e</sup> | + | no |
| Atu0227 | - | tRNA-Leu | 2.15 | - | + | no |
| Atu5167 | *avhB6* | type IV secretion protein | 2.01 | - | + | no |
| Atu2490 | *asd* | aspartate semialdehyde dehydrogenase | 1.93 | - | + | no |
| Atu3318 | - | LuxR family transcriptional regulator | 1.84 | + | + | no |
| Atu1471 | *rluC* | ribosomal large subunit pseudouridine synthase C | 1.84 | - | + | no |
| Atu3755 | *purK* | phosphoribosylaminoimidazole carboxylase ATPase subunit | 1.70 | - | + | no |
| Atu3606 | *ftsE* | cell division ATP-binding protein | 1.63 | + | + | no |
| Atu1791 | - | ABC transporter, membrane spanning protein (sugar) | 1.55 | - | + | no |
| Atu3572 | - | XRE family transcriptional regulator | 1.52 | + | + | yes |
| Atu2217 | - | *hypothetical protein* | 1.52 | - | + | no |
| Atu5119 | *phoB* | two component response regulator | 1.48 | - | - | no |
| Atu3031 | - | *hypothetical protein* | 1.46 | - | - | no |
| Atu1886 | - | *DNA glycosylase* | 1.41 | - | - | no |
| Atu1964 | - | tRNA-Trp | 1.40 | + | + | no |
| Atu1301 | - | *Nudix hydrolase* | 1.39 | - | + | no |
| Atu3610 | - | cation transporter | 1.34 | - | - | no |
| Atu6048 | - | *RNA helicase* | 1.32 | - | + | no |
| Atu1691 | *dgcB* | GGDEF family protein | 1.32 | + | + | no |
| Atu1887 | *exol* | succinoglycan biosynthesis protein | 1.30 | + | + | no |
| Atu1134 | - | *lysyl-phosphatidylglycerol synthase* | 1.30 | - | - | no |
| Atu2665 | - | MarR family transcriptional regulator | 1.27 | - | + | no |
| Atu2204 | - | *hypothetical protein* | 1.25 | - | + | no |
| Atu0540 | - | *hypothetical protein* | 1.25 | - | - | no |
| Atu4856 | - | nucleotidyltransferase | -1.21 | - | - | no |
| Atu4347 | *tae* | type VI secreted effector | -1.23 | (+) Atu4344 | - | no |
| Atu4055 | *exoK* | endo-1,3-1,4-beta-glycanase | -1.28 | (+) Atu4056 | + | yes |
| Atu4345 | *tssD* | type VI secretion needle tube protein | -1.29 | (+) Atu4344 | + | no |
| Atu5091 | *rcdB* | curdlan synthesis protein | -1.32 | + | + | no |
| Atu4357 | - | *transglutaminase-like Cys protease* | -1.34 | - | + | yes |
| Atu0343 | *barA* | two component sensor kinase/response regulator hybrid | -1.37 | - | - | no |
| Atu4053 | *exoA* | succinoglycan biosynthesis protein | -1.42 | (+) Atu4056 | + | yes |
| Atu4056 | *exoH* | succinoglycan biosynthesis protein | -1.42 | + | + | yes |
| Atu3564 | *exsH* | endo-1,3-1,4-beta-glycanase | -1.46 | - | - | no |
| Atu3541 | - | *transglutaminase-like Cys protease* | -1.51 | - | + | no |
| Atu4627 | - | *hypothetical protein* | -1.56 | - | - | no |
| Atu4049 | *exoP* | exopolysaccharide polymerization/transport protein | -1.64 | (+) Atu4056 | + | no |
| Atu1469 | - | hypothetical protein | -1.80 | + | + | no |
| Atu4050 | *exoN* | UTP-glucose-1-phosphate uridylyltransferase | -1.82 | (+) Atu4056 | + | yes |

<sup>a</sup> Gene expression was compared between WT and $\Delta pdhS2$ strains using whole-genome microarrays. Genes were defined as differentially regulated when the following conditions were met: $\log_2 FC \geq 0.50$ OR $\log_2 FC \leq -0.50$ , AND $P < 0.050$ , AND $Q < 0.10$ .
<sup>b</sup> Predicted functions of hypothetical proteins, if available, are italicized.
<sup>c</sup> Intergenic regions 500 nt upstream and 100 nt downstream of start codon for each gene (or operon, if applicable) was scanned for possible CtrA binding sites, as described in the text.
<sup>d</sup> Results were compared with previously published *S. meliloti cbrA* and *divJ* microarrays and *C. crescentus divJ* and *pleC* microarrays, as described in the text.
<sup>e</sup> (+) with Atu number indicates that the putative CtrA box is located in the first gene of a predicted operon

To determine whether any of these 39 genes were putatively regulated by CtrA we scanned a sequence window from 500 bp upstream of the start codon to 100 bp into the coding sequence for plausible CtrA binding sites. CtrA binding sites were defined using the conserved alphaproteobacterial CtrA recognition sequence 5’-TTAANNNNNNGTTAAC-3’ (48, 49). Sequences containing seven or more of the conserved nucleotides in this motif were deemed plausible candidates. Using these parameters 15 differentially transcribed loci are expressed from promoters (some from upstream genes in the operon) with putative CtrA binding sites and are thus putatively directly regulated by CtrA, including *dgcB* and Atu3318, as well as all five of the downregulated succinoglycan biosynthetic genes (Fig. S5; Table 2). We also identified numerous CtrA half-sites containing the sequence 5’-TTAA-3’. In *C. crescentus* CtrA has been shown to bind to such half motifs resulting in transcriptional effects (53). Twenty-eight promoters contained at least one CtrA half-site (Table 2).

To extend our microarray results we measured transcription of translational fusions to β-galactosidase for *dgcB* and Atu3318, in wild-type, Δ*pdhS2*, and Δ*divK* strain backgrounds (Table 1). In both cases β-galactosidase activity increased in the Δ*pdhS2* mutant, corroborating the microarray results. Furthermore, activity decreased from each promoter fusion in the Δ*divK* background supporting inverse regulation by PdhS2 and DivK at these promoters. Finally, to confirm a role for CtrA in transcriptional regulation by PdhS2 and DivK we mutated one CtrA half-site (−126 from start codon, TTAA to AATT) in the *dgcB* promoter region and evaluated transcription from this promoter fused to β-galactosidase as above. Surprisingly, activity from this mutated promoter was diminished in all backgrounds tested (wild-type, Δ*pdhS2*, and Δ*divK*) suggesting that CtrA does interact with this site to influence transcription of *dgcB*, but that this regulation is complex (Table 1). Overall these results are consistent with PdhS2 impacting the motile cell developmental program through CtrA-dependent transcriptional control.

### Evaluation of CtrA protein levels

Turnover of CtrA is known to play a role in its regulatory activity in several related systems. To determine the effect of PdhS2 and DivK on CtrA activity and stability in *A. tumefaciens* we evaluated its steady-state levels and turnover in unsynchronized, stationary phase cultures using western blotting with antibodies raised against *C. crescentus* CtrA. In stationary phase cultures CtrA levels were increased in the Δ*pdhS2* background and decreased in the Δ*divK* background, but these effects were quite modest. This observation is however consistent with PdhS2 and DivK inversely impacting CtrA accumulation or stability (Fig. 4A).

**Figure 4. PdhS2 impacts CtrA abundance but not turnover kinetics.** (A) Steady-state CtrA levels in indicated strains as determined via SDS-PAGE followed by immunoblotting with rabbit anti-CtrA_*Cc*_ primary and IRDye 800CW-conjugated goat anti-rabbit secondary antibodies. (B) Proteolytic turnover of CtrA following translational arrest with chloramphenicol. Aliquots were removed at the indicated time points, lysed, and stored frozen at −20 °C. CtrA levels were then determined via SDS-PAGE followed by immunoblotting, as described for (A) above. Blots were imaged using an LI-COR Odyssey Classic infrared imaging system and band intensities quantified with the Odyssey Classic software. Protein concentrations for each sample were normalized to optical density prior to electrophoresis.

CtrA protein stability was evaluated following treatment of wild-type and Δ*pdhS2* cultures with chloramphenicol to inhibit translation during exponential growth. In wild-type cultures, translation inhibition leads to a decline in CtrA abundance over the course of 2-3 hours, diminishing to roughly 30% of the steady-state levels. Loss of *pdhS2* had no effect on either final steady-state levels of CtrA following inhibition of translation or the observed rate of turnover when compared with wild-type cultures under these conditions (Fig. 4B).

### PdhS2 activity intersects with cyclic-di-GMP pools

In *C. crescentus* DivJ and PleC positively regulate, via phosphorylation, a second response regulator, PleD, as well as DivK (22). In *C. crescentus* and *A. tumefaciens* the *divK* and *pleD* coding sequences form one operon and transcriptional regulation of both genes is linked. Since several PdhS kinases in these systems are predicted to interact with both DivK and PleD we analyzed the effect of loss of PleD activity in the Δ*pdhS2* background. As reported previously, deletion of *A. tumefaciens pleD* alone has only modest effects on swimming motility and adherent biomass (35). Loss of *pleD* in the Δ*pdhS2* background had a minimal effect on swimming motility (Fig. S6). Biofilm formation, however, was reduced by approximately 30%, indicating that PleD contributes to the increased attachment phenotype of the Δ*pdhS2* mutant (Fig. 5A).

**Figure 5. PdhS2 intersects with the activity of multiple diguanylate cyclases.** (A) Biofilm formation was quantified for the wild-type (WT) and indicated mutant strains as described in Figure 2. PleD, DgcA, DgcB have demonstrated *in vivo* diguanylate cyclase enzymatic activity. Thus far conditions under which DgcC is active have yet to be identified. *P* < 0.05 compared to WT (^a^), Δ*pdhS2* (^b^), or corresponding diguanylate cyclase (^c^). (B) The effect on biofilm formation of plasmid-borne expression of wild-type *dgcB* (p-*dgcB*) or a catalytic mutant allele of *dgcB* (p-*dgcB*\*) was evaluated. Expression of each *dgcB* allele was driven by the *P*_*lac*_ promoter. Biofilm formation was evaluated as described in Figure 2. (*) = *P* < 0.05 compared to vector alone.

PleD is a GGDEF motif-containing diguanylate cyclase (DGC), and thus it is likely that the attachment phenotype of the Δ*pdhS2* mutant requires increased levels of cdGMP. Earlier work from our lab identified three additional DGCs that are relevant to attachment and biofilm formation: DgcA, DgcB, and DgcC (52). As seen in wild-type C58, deletion of *dgcA* or *dgcB* in the Δ*pdhS2* background significantly decreased attachment and biofilm formation, whereas loss of *dgcC* did not (Fig. 5A). These data suggest that increased biofilm formation by the Δ*pdhS2* mutant is dependent on cdGMP pools, generated through PleD, DgcA, or DgcB. Loss of *dgcB* largely abolishes the increased biofilm formation of the Δ*pdhS2* mutant. We also compared biofilm formation in Δ*dgcB*Δ*pleD* and Δ*pdhS2*Δ*dgcB*Δ*pleD* mutant backgrounds. In the absence of both DgcB and PleD biofilm formation was enhanced by loss of PdhS2 (Fig. S7). These data suggest that the increased biofilm formation of a *pdhS2* mutant is dependent on a cdGMP pool that is predominantly due to DgcB, but that is also under the cumulative influence of multiple DGC enzymes. Swimming motility was equivalent in either wild-type C58 or Δ*pdhS2* backgrounds in combination with mutations in *pleD*, *dgcA*, *dgcB*, or *dgcC* (Fig. S6 and S7).

Previously we found that mutants lacking either *dgcA*, *dgcB*, or *dgcC* show insignificant differences in total cytoplasmic levels of cdGMP (52). Nonetheless, loss of either *dgcA* or *dgcB*, or mutation of the GGDEF catalytic site of either enzyme, significantly reduced biofilm formation (52), implicating these enzymes in controlling the pool of cdGMP and thereby affecting attachment. We compared cytoplasmic cdGMP levels for wild-type C58, the Δ*pdhS2* mutant, and the Δ*pdhS2*Δ*dgcB* mutant strain and found these levels to be low, with no significant change between mutants (Fig. S8). To verify that the DGC activity is responsible for the increased biofilm formation in the Δ*pdhS2* background we expressed an allele of *dgcB* with a mutation in its GGDEF catalytic motif (GGAAF; *dgcB\**) that abrogates cdGMP formation and that fails to complement a Δ*dgcB* mutant for either cdGMP formation or attachment phenotypes (52). Plasmid-borne expression of wild-type *dgcB* (*P*_*lac*_-*dgcB*) results in a massive increase in attachment and biofilm formation in either the wild-type C58 or Δ*pdhS2* background (Fig. 5B). Expression of *dgcB\** from a *P*_*lac*_ promoter, however, did not increase biofilm formation in either background. In the Δ*pdhS2*Δ*dgcB* mutant background expression of wild-type *dgcB* increased biofilm formation to the same degree seen in the wild-type and Δ*pdhS2* mutant. Expression of the mutant *dgcB*\* allele in the Δ*pdhS2*Δ*dgcB* mutant did appear to modestly affect biofilm formation, although far less than with the wild-type *dgcB* allele. Swimming motility was modestly but significantly reduced when the wild-type *dgcB* allele was provided *in trans* and this was abolished by the *dgcB*\* mutation (Fig. S9). Together our data are consistent with PdhS2-dependent biofilm formation and, to a lesser extent, swimming motility, being mediated at least in part by cdGMP levels.

### Increased attachment in a *pdhS2* mutant requires the UPP polysaccharide

We have previously reported that PleD-stimulated attachment was due to increased levels of the unipolar polysaccharide (UPP) and cellulose (52). In addition to UPP and cellulose, *A. tumefaciens* produces at least three other exopolysaccharides: succinoglycan, cyclic β-1, 2 glucans, β-1, 3 glucan (curdlan), as well as outer membrane associated lipopolysaccharide (LPS) (54). Of these only LPS is essential for *A. tumefaciens* growth (36). The Δ*pdhS2* strain was tested for the impact of each of the non-essential exopolysaccharides for biofilm formation and swimming motility. The *upp* mutation completely abolished attachment in both the wild type and the *pdhS2* mutant (Fig. S10A). The *chvAB* mutant, known to have pleiotropic effects (55), was diminished in adherence overall, but was still elevated by the *pdhS2* mutation. None of the other exopolysaccharide pathways impacted adherence in either background. The decreased swimming phenotype of the *pdhS2* mutant was not significantly altered for any of the exopolysaccharide mutants (Fig. S10B). These results indicate that biofilm formation in the Δ*pdhS2* strain is dependent primarily on UPP production and that the motility phenotype of the Δ*pdhS2* mutant is not dependent on any of the known exopolysaccharides.

## DISCUSSION

### PdhS2 regulates attachment and motility predominantly by its phosphatase activity and through CtrA

Regulation of the developmental program of many alphaproteobacteria centers on the global transcriptional regulator CtrA (43, 49, 56, 57). CtrA activity is controlled, indirectly, through a series of phosphotransfer reactions dependent on one or more PdhS-type histidine kinases. Here we show that PdhS2, one of at least four PdhS family kinases from *A. tumefaciens*, regulates motility and attachment at least in part through fine-tuning of CtrA activity, thereby impacting the CtrA regulon. We demonstrate that null mutations of the single domain response regulator *divK* are epistatic to *pdhS2* mutations in *A. tumefaciens*. Mutation of specific DGCs also reverse the phenotypes of *pdhS2* mutants, suggesting that elevated attachment and decreased motility are mediated through cdGMP pools. The phosphatase activity of PdhS2 is predominantly responsible for its function during laboratory culture growth. It is worth noting however, that strains bearing the phosphatase-null and kinase-null alleles of PdhS2 do not phenocopy one another nor the Δ*pdhS2* mutant strains, supporting distinct roles for both enzymatic activities. This is in contrast to mutant alleles of *pleC* in *C. crescentus* (46, 58, 59). Fine-scale determination of the timing, specificity, and regulation of both phosphatase and kinase activities await further experimentation.

### Indirect regulation of CtrA by PdhS2

Our data support altered CtrA activity as responsible for many of the *pdhS2*-dependent transcriptional responses, and through these motility and attachment. A significant fraction of the differentially expressed genes had presumptive CtrA binding sites in their upstream regions. Several of the genes examined directly were dysregulated in both *pdhS2* and *divK* mutants, but usually in the opposite directions. We predict that both DivK and PdhS2 function through affecting the phosphorylation state of CtrA. In *A. tumefaciens* CtrA activates certain promoters (*e.g.* succinoglycan synthetic genes and *ctrA*) while repressing others (*e.g. dgcB*, Atu3318, and *pdhS1*). Although the *pdhS2* and *divK* mutations do slightly change steady state levels of CtrA, our findings suggest that changes in CtrA stability do not explain the resulting phenotypes. We posit that the phosphorylation state of CtrA is the dominant mechanism by which its activity is affected in these mutants. There are several examples in *C. crescentus* where altered CtrA activity is observed without significant changes in CtrA abundance (36, 60, 61). There are also several examples in both *C. crescentus* and *S. meliloti* where altered PdhS kinase activity results in both perturbed activity and altered abundance of CtrA (42, 62). The subtle effects on CtrA abundance coupled with the modest, but significant, effects we observed for several CtrA-dependent promoters, suggest that PdhS2 regulation of CtrA activity may be restricted to a very tight window during the cell cycle. Alternatively, the complement of remaining PdhS kinases may buffer CtrA activity in the absence of *pdhS2*.

### PdhS2 influences cdGMP-dependent phenotypes

The increased biofilm formation and diminished motility in a *pdhS2* mutant is most similar to the inverse regulation frequently observed for increasing internal pools of cdGMP (63). Indeed, our data demonstrate a strong dependence on specific diguanylate cyclases for mediating the Δ*pdhS2* hyperadherent phenotype, and transcription analysis demonstrates that *dgcB* expression is elevated in this mutant. Although measurements of the cytoplasmic cdGMP levels suggest that they remain low overall in the *pdhS2* mutant, it is clear that mutations in *dgcB* strongly reverse the effects of the *pdhS2* mutation on attachment and that, to a lesser extent, mutations in *dgcA* and *pleD* can diminish them. This suggests that increased cdGMP synthesis via these enzymes may impart the effect on UPP-dependent attachment. Although DgcB seems to have the dominant effect, it is plausible that PdhS2 may affect PleD DGC activity through the phosphorylation state of its receiver domain, similar to other PleD homologues (22, 64, 65). In *A. tumefaciens* PleD has only modest effects on motility and attachment, and does not significantly contribute to cell cycle control (35). Neither DgcB nor DgcA is a response regulator, and it is more likely that at least for *dgcB,* its elevated expression is the mechanism through which CtrA functions. Recent studies have revealed that the *C. crescentus* orthologue of *A. tumefaciens* DgcB regulates holdfast synthesis in response to changes in flagellar rotation (66). Thus, changes in *dgcB* levels due to mutation of *pdhS2* may be also be impacting the motile to sessile transition in *A. tumefaciens*.

The phenotypes regulated by PdhS2 mirror those regulated by the master motility regulators VisR and VisN (52). Loss of either *visN* or *visR* results in abolishment of motility and a dramatic increase in attachment that is dependent on cdGMP production and the UPP adhesin. However, the motility defect in *visNR* mutants is predominantly transcriptional, as expression of all of the flagellar genes is dramatically decreased. In contrast, none of the flagellar genes are differentially regulated in the *pdhS2* mutant as measured in our microarray data, and flagella are assembled but decreased in activity. The increased attachment in *visNR* mutants is however due to elevated *dgcB* expression and also requires *dgcA*, through increased cdGMP and elevation of UPP and cellulose biosynthesis (52). Interestingly, other target genes that are derepressed in *pdhS2* mutants are also among the small fraction of the genes that are increased in a *visR* mutant, such as Atu3188. Given the presence of CtrA boxes in their upstream sequences this may suggest a common underlying mechanism. Interestingly, mutation of *dgcB* or the other DGCs does not enhance the dramatically impeded motility of the *pdhS2* mutant. This suggests that the loss of motility in the *pdhS2* mutant is not primarily due to elevated cdGMP levels. In many systems, CtrA directly regulates motility, often through flagellar gene expression (67, 68). In fact, plasmid-borne expression of the *A. tumefaciens* CtrA in a *ctrA* null mutant of the marine alphaproteobacterium *Ruegeria* sp. KLH11 (*ctrA* is not essential in this taxon), effectively reverses its non-motile phenotype (69), indicative of its positive impact on motility on this bacterium. It seems likely that the non-motile phenotype of the *A. tumefaciens pdhS2* mutant likewise reflects a decrease in active CtrA.

Interestingly, as shown for *C. crescentus* and *A. tumefaciens*, elevated cdGMP allosterically switches the bifunctional hybrid histidine kinase CckA from kinase to phosphatase mode, thereby downregulating CtrA phosphorylation and DNA-binding activity (30, 31). Thus effects on local cdGMP levels can feed back on the CtrA pathway, reinforcing decreases in CtrA~P that would be coincident with increased cdGMP.

### Segregation of antagonistic signaling activity promotes asymmetric development

The asymmetric division of *A. tumefaciens* and other alphaproteobacteria, producing two genetically identical but phenotypically distinct daughter cells, requires well-coordinated regulation of two developmental programs. The mother cell remains in a terminally differentiated state, proceeding through distinct synthesis (S) and growth (G1/G2) phases of the cell cycle (54, 56). During G1/G2 phase the cell elongates into a predivisional cell and establishes a functional asymmetry between its two cellular poles by differential localization of antagonistic homologues of the PdhS kinases. At least one PdhS kinase localizes to the old pole; DivJ in *C. crescentus*, DivJ and PdhS1 in *A. tumefaciens* (Fig. 1A and 3B), PdhS in *B. abortus*, and CbrA in *S. meliloti*, (18, 41, 47). From this position these kinases can act to phosphorylate targets such as DivK and PleD, indirectly inactivating CtrA (as reported for *C. crescentus*). At the opposite pole at least one PdhS kinase, PleC in *C. crescentus* and PdhS2 in *A. tumefaciens* (Fig. 1A and 3A), localizes and acts primarily through its phosphatase activity to dephosphorylate targets, ultimately promoting CtrA stability and activity. Upon cytokinesis, then, the motile daughter cell is released in a G1/G2 growth phase with high levels of CtrA activity establishing a distinct transcriptional program and limiting DNA replication.

Our data are consistent with PdhS2 acting in the motile daughter cell to prevent premature activation of cell attachment processes, as well as to promote motility. PdhS2 dynamically localizes to the new pole of *A. tumefaciens* cells following cytokinesis while DivJ, another PdhS-type kinase, localizes to the old pole of each cell. We propose that together the antagonistic activities of DivJ and PdhS2 (and perhaps additional PdhS homologues), coupled with their distinct localization patterns, generate a spatiotemporal gradient of phospho-CtrA, thus differentially regulating the developmental program of *A. tumefaciens* (Fig. 1A). Localized synthesis and degradation of cdGMP contributes to this regulatory gradient. Prior to and after cytokinesis daughter cells would have PdhS2 at their flagellar pole, reinforcing the CtrA pathway, increasing CtrA~P, promoting motility and preventing adhesive processes. In contrast, at the mother cell old pole after PdhS2 delocalization, DivJ kinase activity would dominate, the CtrA pathway would be inhibited, and lower CtrA~P levels would promote DNA replication, maintaining a sessile, non-motile state. Computational models of asymmetric cell development in *C. crescentus* support this notion, with the important *caveat* that phospho-DivK may not be distributed in a gradient but rather locally restricted (23, 25, 70).

### PdhS2 may regulate CtrA activity via an alternate route

The recognized architecture of the DivK-CtrA regulatory pathway in several alphaproteobacteria, coupled with our data demonstrating a genetic interaction between *divK* and *pdhS2* in *A. tumefaciens*, are consistent with PdhS2, primarily through its phosphatase activity, decreasing DivK phosphorylation, similar to what is predicted for the other PdhS-type kinase PleC (Fig. 6, Model A). The *pdhS2* mutant phenotype is however in stark contrast to the other non-essential *A. tumefaciens* PdhS-type mutants and the *divK* mutant, which all cause cell branching (35). How does PdhS2 regulate the same pathway so differently from the PdhS-type proteins? Possibly, spatial restriction of PdhS2 activity to the new poles of mother cells, that rapidly transition to become the old pole of newly formed daughter cells, imparts PdhS2 control of motility and attachment processes, without strongly influencing the budding process *per se*. Alternatively, PdhS2 may act via a different mechanism to influence CtrA activity.

**Figure 6. An alternative model for PdhS2 regulation of CtrA activity.** Our data are consistent with PdhS2 intersecting the DivK-CtrA regulatory pathway at one of two points. (Pathway A) Canonical genetic model with PdhS2 interacting with DivK. The phosphorylation status of DivK then modulates CtrA activity through the CckA-ChpT-CtrA axis. (Pathway B) DivK-independent model of CtrA regulation by PdhS2 through an unidentified response regulator, RR-X. Both routes to regulation of CtrA activity ultimately affect the phosphorylation status of CtrA, affecting occupancy at CtrA-regulated promoters, and finally leading to inverse regulation of attachment (primarily through cdGMP pools) and separately motility. Regulatory proteins: Blue text; histidine kinases; orange text, histidine phosphotransferase (Hpt); green text, response regulators. RR-X indicates a putative response regulator, yet to be identified.

An interesting possibility is that PdhS2 and DivK may work in parallel rather than in series to impact CtrA activity and its target genes (Fig. 6, Model B). The apparent epistasis of the *divK* mutation over the *pdhS2* mutation could result from the unfettered activity of CckA in the *divK* mutant, which titrates the impact of the *pdhS2* mutation. Our results in which expression of wild-type and kinase-locked CckA (CckA^Y67D^) alleles in the Δ*pdhS2* mutant only modestly impact its mutant phenotypes, support this proposal (Fig. S3 and S4). The CckA^Y67D^ mutant was isolated as a spontaneous suppressor of the swimming deficiency of a *pleC* mutant (35). Plasmid-borne ectopic expression of the CckA^Y67D^ effectively reversed *pleC* phenotypes, in contrast to the observation that it does not suppress *pdhS2* mutant phenotypes (Fig. S3 and S4). This suggests that PdhS2 does not act similarly to PleC to inhibit DivK phosphorylation. A plausible explanation is that PdhS2 control of CtrA activity is independent of DivK and CckA (Fig. 6, Model B). Although uninhibited CckA kinase activity in a *divK* mutant can overcome the effect of the *pdhS2* mutation, perhaps the kinase-locked CckA^Y67D^ allele is insufficiently active to do so.

In this model, PdhS2 intercepts the DivK-CtrA signaling axis at a node downstream or independent of CckA (Fig. 6). Our findings reveal that the phosphatase activity of PdhS2 is dominant in its impact on CtrA-dependent targets, suggesting that it dephosphorylates a response regulator that itself is inhibitory to CtrA activity. A response regulator that inhibits CtrA directly is CpdR (Fig. 6), which in *C. crescentus* stimulates proteolytic turnover of CtrA (71). *A. tumefaciens* has two CpdR homologues, CpdR1 and CpdR2 (Atu3883 and Atu3603, respectively). *S. meliloti* likewise maintains two CpdR homologues one of which, CpdR1, impacts developmental phenotypes and CtrA activity (62, 72). However, current models suggest that for CpdR in *C. crescentus* and CpdR1 in *S. meliloti*, phosphorylation through ChpT decreases its ability to drive CtrA degradation. Therefore, in *A. tumefaciens*, if PdhS2 acts on the pathway through its phosphatase activity, dephosphorylation of CpdR1 would increase its inhibitory capacity for CtrA. Mutation of *pdhS2* would then be predicted to lead to more CtrA~P available to inhibit *dgcB* expression and stimulate motility, generating the opposite of the non-motile, hyperadherent phenotypes we observe. It is formally possible that in *A. tumefaciens* CpdR functions differently than its orthologues in *C. crescentus* or *S. meliloti,* or that the PdhS2 target is a different response regulator.

A plausible alternative target for PdhS2 is CpdR2 (Atu3603). The CpdR2 response regulator of *S. meliloti* does not impact developmental phenotypes (77), consistent with our own observations for *cpdR2* mutants of *A. tumefaciens* (Heindl et al. unpublished results). If not CpdR2, the target could be a response regulator (RR-X) that is thus far unrecognized to function in CtrA control (Fig. 6). In either case, one would predict that the phosphorylated form of the response regulator is active for CtrA inhibition, and that dephosphorylation by PdhS2 diminishes its inhibitory activity on CtrA. Conversely, *pdhS2* mutants promote the strongly activated form of this response regulator, inhibiting CtrA, derepressing *dgcB* expression and limiting motility, and in turn driving cdGMP-responsive attachment. We have previously proposed additional direct targets for the PdhS-type kinase DivJ as a means to explain the contradiction between the essentiality of *divJ* and the non-essentiality of *divK* in *A. tumefaciens* (35). In that work we also noted the possibility of DivJ directly targeting CtrA as observed *in vitro* for *C. crescentus* DivJ. It is thus possible that both DivJ and PdhS2 in *A. tumefaciens* act downstream of CckA, perhaps both through RR-X (Fig. 6), more directly influencing CtrA (73).These models are currently being tested, but are more challenging due to the essentiality of many of the regulatory components in this domain of the pathway for *A. tumefaciens*, including DivJ itself.

## MATERIALS AND METHODS

### Strains and plasmids

Bacterial strains, plasmids, and oligonucleotides used in these studies are listed in Tables S1 through S3. *A. tumefaciens* was routinely cultivated at 28°C in AT minimal medium plus 1% (w/v) glucose as a carbon source and 15 mM (NH_4_)_2_SO_4_ as a nitrogen source (ATGN), without exogenous FeSO_4_ (74, 75). For biofilm assays 22 μM FeSO_4_ was included in the media. *E. coli* was routinely cultivated at 37°C in lysogeny broth (LB). Antibiotics were used at the following concentrations (*A. tumefaciens*/*E. coli*): ampicillin (100/100 μg·mL^-1^), kanamycin (150/25 μg·mL^-1^), gentamicin (150/30 μg·mL^-1^), spectinomycin (300/100 μg·mL^-1^), and tetracycline (4/10 μg·mL^-1^).

Non-polar, markerless deletion of *pdhS2* (Atu1888) in all genetic backgrounds used in this work was accomplished using <u>s</u>plicing by <u>o</u>verlap <u>e</u>xtension (SOE) polymerase chain reaction (PCR) followed by homologous recombination, as described (35). Suicide plasmid pJEH040 carries an approximately 1 kb SOE deletion fragment of *pdhS2* on a pNPTS138 vector backbone. pNPTS138 is a ColE1 plasmid and as such is unable to replicate in *A. tumefaciens*. pJEH040 was delivered to recipient strains by either transformation or conjugation followed by selection on ATGN plates supplemented with 300 μg·mL^-1^ Km, selecting for *A. tumefaciens* cells in which pJEH040 had integrated at the chromosomal *pdhS2* locus by homologous recombination. Recombinants were then grown overnight at 28°C in ATGN in the absence of Km and plated the following day onto ATSN (ATGN with sucrose substituted for glucose) agar plates to select for sucrose resistant (Suc^R^) allelic replacement candidates. After three days’ growth at 28°C colonies were patched in parallel onto ATGN Km and ATSN plates. Km^S^ Suc^R^ recombinants were then tested for the targeted deletion by diagnostic PCR using primers external to the *pdhS2* locus (JEH100 and JEH113) as well as internal primers (JEH85 and JEH87). Candidate colonies were further streak purified and verified a second time by diagnostic PCR before being used in downstream assays. Non-polar, markerless deletion of *dgcB* (Atu1691) in the Δ*pleD* and Δ*pdhS2*Δ*pleD* genetic backgrounds was achieved using the above strategy with the pNPTS138 derivative pJX802.

Site-directed mutagenesis of *pdhS2* was achieved using mutagenic primer pairs JEH245/JEH246 (for generating the His271Ala allele, H271A) or JEH261/JEH262 (for generating the Thr275Ala allele, T275A). Plasmid pJEH021 carrying the wild-type *pdhS2* sequence was amplified by PCR using the above primer pairs. Following amplification, reaction mixtures were treated with *Dpn*I restriction endonuclease to remove template plasmid and then transformed into TOP10 F’ *E. coli* competent cells. Purified plasmids from each transformation were sequenced and those containing the desired mutations, pJEH091 for His271Ala and pJEH099 for Thr275Ala, were selected for sub-cloning. pJEH091 and pJEH099 were digested with *Nde*I and *Nhe*I followed by gel electrophoresis and purification of the resulting insert. Inserts were ligated into similarly digested pSRKGm and transformed into competent *E. coli* TOP10 F’ cells. Purified plasmids from each transformation were sequenced to verify their identity. The resulting plasmids, pJEH092 (H271A) and pJEH102 (T275A), were used to transform *A. tumefaciens*. To generate a PdhS2 allele carrying both H271A and T275A mutations the same steps were followed as above using plasmid pJEH091 as template with mutagenic primers JEH261/JEH262. Site-directed mutagenesis of the second CtrA half-site (5’-TTAA-3’ → 5’-AATT-3’) located 126 bp upstream of the start codon was performed as above using mutagenic primer pairs USP073/USP074 and plasmid pJX162 as template.

Translational fusions of full-length wild-type PdhS2 and DivJ to GFP were constructed as follows. *pdhS2* and *divJ*, each lacking a stop codon were amplified by PCR using primer pairs JEH65/JEH146 (*pdhS2*) and JEH147/JEH148 (*divJ*) with *A. tumefaciens* strain C58 genomic DNA as template. Primer design for these amplifications included 5’ *Nde*I and 3’ *Nhe*I restriction sites. The *gfpmut3* gene including a 5’ *Nhe*I site and a 3’ *Kpn*I site was amplified using primer pair JEH149/JEH150 and pJZ383 as template. Amplicons were gel purified, ligated into pGEM-T Easy, transformed into competent TOP10 F’ *E. coli*, and eventually sequenced. The resulting plasmids, pJEH052 (*pdhS2*), pJEH053 (*gfpmut3*), and pJEH054 (*divJ*) were digested with either *Nde*I and *Nhe*I (pJEH052 and pJEH054) or *Nhe*I and *Kpn*I (pJEH053). Inserts were gel purified and used in a three-component ligation with *Nde*I/*Kpn*I-digested pSRKGm generating pJEH060 (PdhS2-GFP) and pJEH078 (DivJ-GFP). Sequenced plasmids were used to transform *A. tumefaciens*.

Reporter gene fusion constructs included predicted promoter regions from between 200 bp and 400 bp upstream of the indicated gene through the start codon. Each upsream region was amplified by PCR using the primers listed in Table S3 using *A. tumefaciens* genomic DNA as template. Amplicons were gel purified, ligated into pGEM-T Easy, transformed into competent TOP10 F’ *E. coli*, and eventually sequenced. The resulting plasmids, pJEH113 (*ccrM*, Atu0794, promoter), pJEH115 (*ctrA*, Atu2434, promoter), and pJEH119 (*pdhS1*, Atu0614, promoter) were digested with either *Kpn*I and *Hin*DIII (pJEH113 and pJEH119) or *Kpn*I and *Pst*I (pJEH115). Inserts were gel purified and ligated with similarly cleaved pRA301 containing a promoterless *E. coli lacZ* gene without its own ribosome binding site. The resulting constructs (pJEH121, pJEH122, and pJEH124) carry *lacZ* translationally fused to the start codon for each gene with transcription and translation driven by the fused upstream region. pJEH121, pJEH122, and pJEH124 were used to transform *A. tumefaciens* for subsequent beta-galactosidase assays.

### Static biofilm assays

Overnight cultures in ATGN were sub-cultured in fresh ATGN to an optical density at 600 nm (OD_600_) of 0.1 and grown with aeration at 28°C until an OD_600_ of 0.25-0.6. Cultures were diluted to OD_600_ of 0.05 and 3 mL were inoculated into each of four wells in a 12-well plate. A single coverslip was placed vertically into each well to submerge approximately half of each coverslip. Plates were incubated in a humidified chamber at 28° for 48 h. Coverslips were removed from each well, rinsed with water, and adherent biomass stained by 5 min immersion in a 0.1% (w/v) crystal violet solution. Adsorbed crystal violet was solubilized by immersion in 1 mL 33% acetic acid and the absorbance of this solution determined at 600 nm (A_600_) on a Synergy HT multi-detection microplate reader (Bio-Tek). Culture density for each sample was also determined by measuring the OD_600_ of each culture. Data are typically presented as A_600_/OD_600_ ratios normalized to values obtained for the wild-type strain within each experiment. ATGN was supplemented with antibiotics and 250 μM IPTG as appropriate. Final inoculations also included supplemental FeSO_4_ (22 μM). Each mutant was evaluated in three independent experiments each of which contained three technical replicates.

### Motility assays

Wet mounts of exponentially growing cultures were observed under brightfield optics using a Zeiss Axioskop 40 equipped with an AxioCam MRm monochrome digital camera. Swim plates containing 0.3% agarose in ATGN, supplemented with 1 mM IPTG and antibiotics when appropriate, were inoculated with a single colony of the indicated strain at a central point and incubated for 7 days at 28°C. Swim ring diameters were measured daily for seven days. Each experimental condition was tested in three independent experiments containing three technical replicates.

### Microscopy

Cell morphology and localization of PdhS2-GFP and DivJ-GFP was evaluated using a Nikon E800 fluorescence microscope equipped with a Photometrics Cascade cooled CCD camera. Overnight cultures were grown in ATGN with gentamicin and 250 μM IPTG. The following day each strain was sub-cultured to OD_600_ 0.1 and then grown at 28°C with aeration until ~OD_600_ 0.5-0.8. The culture (0.5 μl) was transferred to a 1% ATGN/agarose pad on a clean glass slide and a clean 22 × 22 mm number 1.5 glass coverslip placed on top. Images were acquired using a 100X oil immersion objective and phase contrast optics or epifluorescence with a FITC-HYQ filter set (Nikon; excitation filter = 480/40 nm, dichromatic mirror = 505 nm, absorption filter = 535/50 nm). Time-lapse microscopy utilized a Nikon Ti-E inverted fluorescence microscope with a Plan Apo 60X/1.40 oil Ph3 DM objective, a DAPI/FITC/Cy3/Cy5 filter cube, an Andor iXon3 885 EMCCD camera, and a Lumencor Spectra X solid state light engine at 20% power. For time-lapse imaging agarose pads included 250 μM IPTG and coverslips were attached to the glass slide using a gas-permeable 1:1:1 mixture of Vaseline, lanolin, and paraffin. Phase and fluorescence images were captured every 20 min for 8 h using a 60 ms (phase) or 2 s (fluorescence) exposure. Images were analyzed using ImageJ (76–78).

### Transcriptional profiling

Whole-genome transcriptional profiling using custom 60-mer oligonucleotide microarrays was performed essentially as previously described (79). Arrays were produced by Agilent Technologies, and consist of 8455 features that represent 5338 predicted protein-encoding open reading frames, tRNA and rRNA encoding genes, and 2,983 duplicate spots. Cultures of wild-type or the Δ*pdhS2* mutant strain of *A. tumefaciens* strain C58 were grown overnight in ATGN to full turbidity and then sub-cultured 1:150 into fresh ATGN for a second overnight growth. The following morning a volume equivalent to 11 ml of OD_600_ 0.6 was prepared for RNA extraction using RNAprotect Bacteria Reagent (QIAGEN, Germantown, MD) following the manufacturer’s protocol. RNA was extracted from these samples using QIAGEN RNA midipreps (QIAGEN, Germantown, MD) following the manufacturer’s protocol. DNA contamination was removed by DNase digestion using the TURBO DNA-free kit (Ambion, Austin, TX) with the incubation time extended to two hours. First strand cDNA synthesis was performed using Invitrogen SuperScript Indirect Labeling Kit, and cDNA was purified on Qiagen QIAQuick columns. cDNA was labeled with AlexaFluor 555 and 647 dyes using Invitrogen SuperScript cDNA Labeling Kit, and repurified on QIAQuick columns. cDNA was quantified on a NanoDrop spectrophotometer. Hybridization reactions were performed using Agilent in situ Hybridization Kit Plus, boiled for 5 min at 95°C, applied to the printed arrays, and hybridized overnight at 65°C. Hybridized arrays were washed with Agilent Wash Solutions 1 and 2, rinsed with acetonitrile, and incubated in Agilent Stabilization and Drying Solution immediately prior to scanning the arrays. Three independent biological replicates were performed, with one dye swap. Hybridized arrays were scanned on a GenePix Scanner 4200 in the Center for Genomics and Bioinformatics (CGB) at Indiana University. GenePix software was used to define the borders of hybridized spots, subtract background, measure dye intensity at each spot, and calculate the ratio of dye intensities for each spot. Analysis of the scanned images was conducted using the LIMMA package in R/Bioconductor. Background correction of the data was performed using the minimum method (80, 81). The data was normalized within arrays with the LOESS method, and between arrays with the quantile method. Statistical analysis was performed using linear model fitting and empirical Bayesian analysis by least squares. Genes with significant *P* values (≤ 0.05) and with log_2_ ratios of ≥ 0.50 or ≤ −0.50 (representing a fold-change of ± 1.4) are reported here. Expression data have been deposited in the Gene Expression Omnibus (GEO) database at the National Center for Biotechnology Information (NCBI) under accession number GSE71267 (82).

β-galactosidase activity was measured using a modified protocol of Miller (83). Cultures carrying transcriptional reporter plasmids were grown overnight in ATGN and sub-cultured the following morning to OD_600_ 0.15. Diluted cultures were grown at 28°C with aeration until reaching mid-exponential growth. Between 100 and 300 μL of exponential phase culture was mixed with Z buffer (60 mM Na_2_HPO_4_, 40 mM NaH_2_PO_4_, 10 mM KCl, 1 mM MgSO_4_, pH 7.0) to a final volume of 1 mL (volume of culture = f) plus two drops 0.05% sodium dodecyl sulfate and 3 drops CHCl_3_. The amount of culture volume used was calibrated to generate reaction times between 15 minutes and two hours for cultures with activity. 0.1 mL of a 4 mg·mL^-1^ solution in Z buffer of the colorimetric substrate 2-nitrophenyl β-D-galactopyranoside (ONPG) was added and the time (t) required for the solution to turn yellow was recorded. The reaction was stopped by addition of 1 M Na_2_CO_3_ and the absorbance at 420 nm (A_420_) of each solution was measured. Promoter activity is expressed in Miller units (MUs = [1000 × A_420_nm]/[OD_600_nm × t × f]). Each mutant was tested in three independent experiments containing five technical replicates.

### Protein stability assays

Steady-state levels of CtrA were determined from stationary phase cultures of wild-type, Δ*divK*, and Δ*pdhS2* strains of *A. tumefaciens*. Overnight cultures of each strain were grown in TY broth at 28°C with aeration to an OD_600_ > 1. Two 1 mL aliquots were removed from each culture, pelleted by centrifugation (13,200 × *g*, 2 min) and supernatants discarded. One of the resulting pellets was resuspended on ice in 50 μL 100 mM Tris·HCl, pH 6.8, followed by 50 μL 2X SDS-PAGE loading buffer (65.8 mM Tris·HCl, pH 6.8, 26.3% (v/v) glycerol, 2.1% (w/v) sodium dodecyl sulfate (SDS), 0.01% (w/v) bromophenol blue), then stored frozen at −20°C. The second pellet was resuspended in 100 μL 1X protein assay buffer (32.9 mM Tris·HCl, pH 6.8, 1%SDS), boiled 10 minutes, and used for protein concentration determination using the Pierce BCA Protein Assay Kit (Thermo Fisher Scientific), per manufacturer’s instructions. Frozen resuspended pellets were thawed on ice and β-mercaptoethanol added to a final concentration of 5% prior to electrophoresis. Samples were normalized for protein concentration and separated on a 12.5% SDS-polyacrylamide gel. Following electrophoresis proteins were transferred to Immobilon-FL polyvinyl difluoride membranes (EMD Millipore). Membranes were rinsed in 1X Tris-buffered saline (TBS; 50 mM Tris·HCl, pH 7.5, 150 mM NaCl) solution and air dried. Membranes were wetted with MeOH and incubated in blocking buffer (1X TBS, 5% non-fat dairy milk [NFDM]) for 1 h at room temperature and then incubated overnight at 4°C with primary antibody (1:5000 dilution of rabbit anti-CtrA from *C. crescentus,* anti-CtrA_*Cc*_, in 1X TBS/5% NFDM/0.2% Tween 20). The following day membranes were rinsed thoroughly with 1X TBS/0.1% Tween 20 and incubated 1 h at room temperature with secondary antibody (1:20,000 dilution of IRDye 800CW-conjugated goat anti-rabbit antibody (LI-COR) in 1X TBS/5% NFDM/0.2% Tween 20/0.01% SDS). Membranes were rinsed thoroughly with 1X TBS/0.1% Tween 20 followed by 1X TBS alone and air dried in the dark. The resulting blot was imaged using a LI-COR Odyssey Classic infrared imaging system. Band intensities were quantified using the Odyssey Classic software.

Proteolytic turnover of CtrA was evaluated using translational shut-off assays. Overnight cultures were grown in TY broth at 28°C with aeration. The following day each strain was sub-cultured in fresh TY broth to an OD_600_ 0.05 and incubated at 28°C with aeration. To inhibit protein synthesis 90 μg·mL^-1^ chloramphenicol was added to each culture at OD_600_ 0.5. Starting at the time of chloramphenicol addition 5 mL aliquots were removed every 30 min for 3 h. Each aliquot was pelleted by centrifugation (5000 × *g*, 10 min). Cleared supernatants were discarded and pellets resuspended to an OD_600_ 10.0 in Tris-Cl, (10 mM, pH 8.0). Resuspended pellets were mixed with 2X SDS-PAGE loading buffer and stored frozen at −20°C. Levels of CtrA were determined by SDS-PAGE and Western blotting as described above. Band intensities were quantified using the Odyssey Classic software and normalized to the band intensity of CtrA from the wild-type background at t = 0 min.

### Global cdGMP measurement

Measurement of cdGMP levels was performed by liquid chromatography, tandem mass spectrometry (LC-MS/MS) on a Quattro Premier XE mass spectrometer coupled with an Acquity Ultra Performance LC system (Waters Corporation), essentially as previously described (84). Concentrations of cdGMP in cell samples were compared to chemically synthesized cdGMP (Axxora) dissolved in water at concentrations of 250, 125, 62.5, 31.2, 15.6, 7.8, 3.9, and 1.9 nM to generate a calibration curve. *A. tumefaciens* derivatives were grown in ATGN overnight at 28°C to stationary phase. Culture densities were normalized after collecting cells by centrifugation and then resuspension in the appropriate volume of ATGN. Cultures were then pelleted by centrifugation and resuspended in ice-cold 250 μL extraction buffer (methanol:acetonitrile:water, 40:40:20 + 0.1 N formic acid) and incubated for 30 min at −20°C. Resuspensions were transferred to microcentrifuge tubes and pelleted (13,000 × rpm, 5 min). 200 μL of the resulting supernatant was neutralized with 8 μL 15% NH_4_HCO_3_. Neutralized samples were stored at −20°C. Prior to mass spectrometric analysis, samples were vacuum centrifuged to remove extraction buffer and resuspended in an equal volume of deionized water.

## ACKNOWLEDGEMENTS

This project was supported by National Institutes of Health (NIH) grants GM080546 and GM120337 (C.F.) and GM109259 (C.M.W.). J.E.H. was supported by a Ruth L. Kirschstein National Research Service Award (1 F32 GM100601) from the NIH and the Milton Lev Memorial Faculty Research Fund from University of the Sciences in Philadelphia. The authors thank Peter Chien (U. Mass. Amherst) for generously providing the CtrA_Cc_ antibody, and the Brun laboratory for access to and training on the Nikon E800 microscope.

## SUPPLEMENTARY MATERIAL

**Figure S1.**
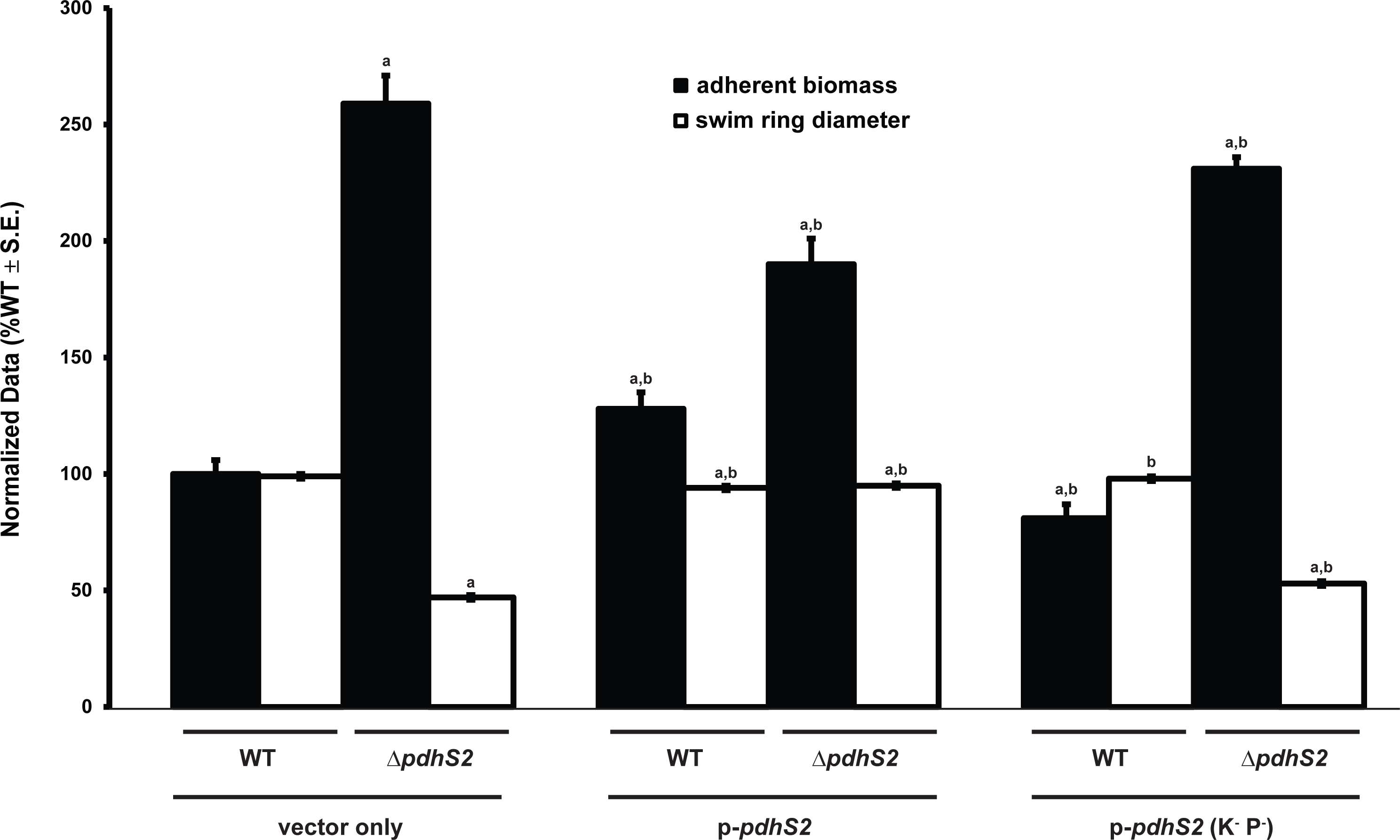
A combined kinase- and phosphatase-null PdhS2 mutant allele has little effect on biofilm formation or swimming motility. The ability of plasmid-borne expression of a kinase- and phosphatase-null allele of *pdhS2* (p-*pdhS2* (K^-^P^-^)) to complement the Δ*pdhS2* phenotypes was compared against the wild-type *pdhS2* allele (p-*pdhS2*). Biofilm formation (black bars) and swimming motility (white bars) were evaluated as in Figure 2. (^a^) = *P* < 0.05 compared to the wild-type background with vector only; (^b^) = *P* < 0.05 compared to the Δ*pdhS2* background with vector only. Statistical significance was determined using Student’s *t* test.

**Figure S2.**
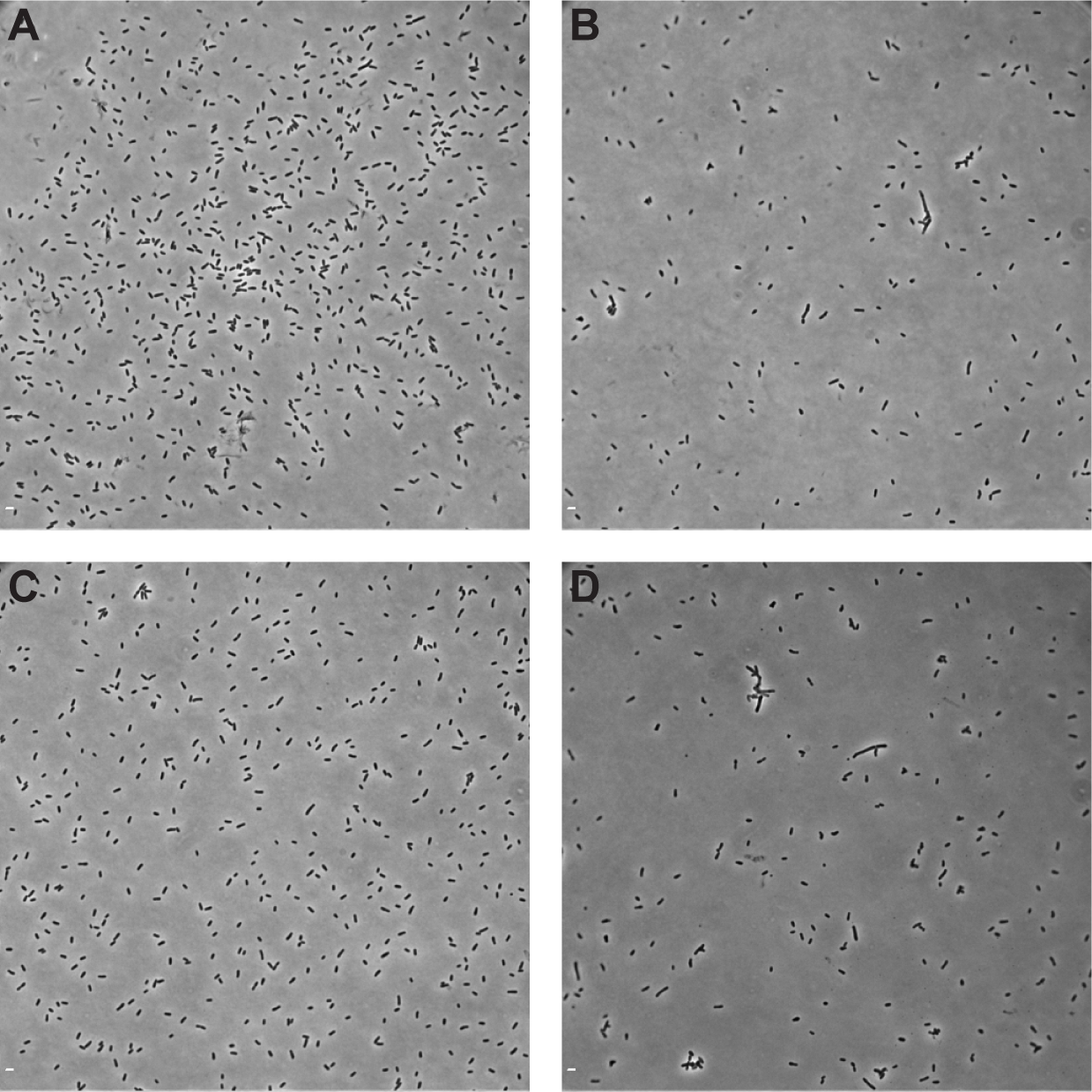
Morphology of WT, Δ*divK*, Δ*pdhS2*, and Δ*divK* Δ*pdhS2* strains. Strains were grown to exponential phase in ATGN. Aliquots of cells were placed on top of an ATGN/1% agarose pad and imaged using phase contrast microscopy. (A), WT; (B) Δ*divK*; (C) Δ*pdhS2*; (D) Δ*divK* Δ*pdhS2*. Representative images are shown. Scale bar = 2 μm.

**Figure S3.**
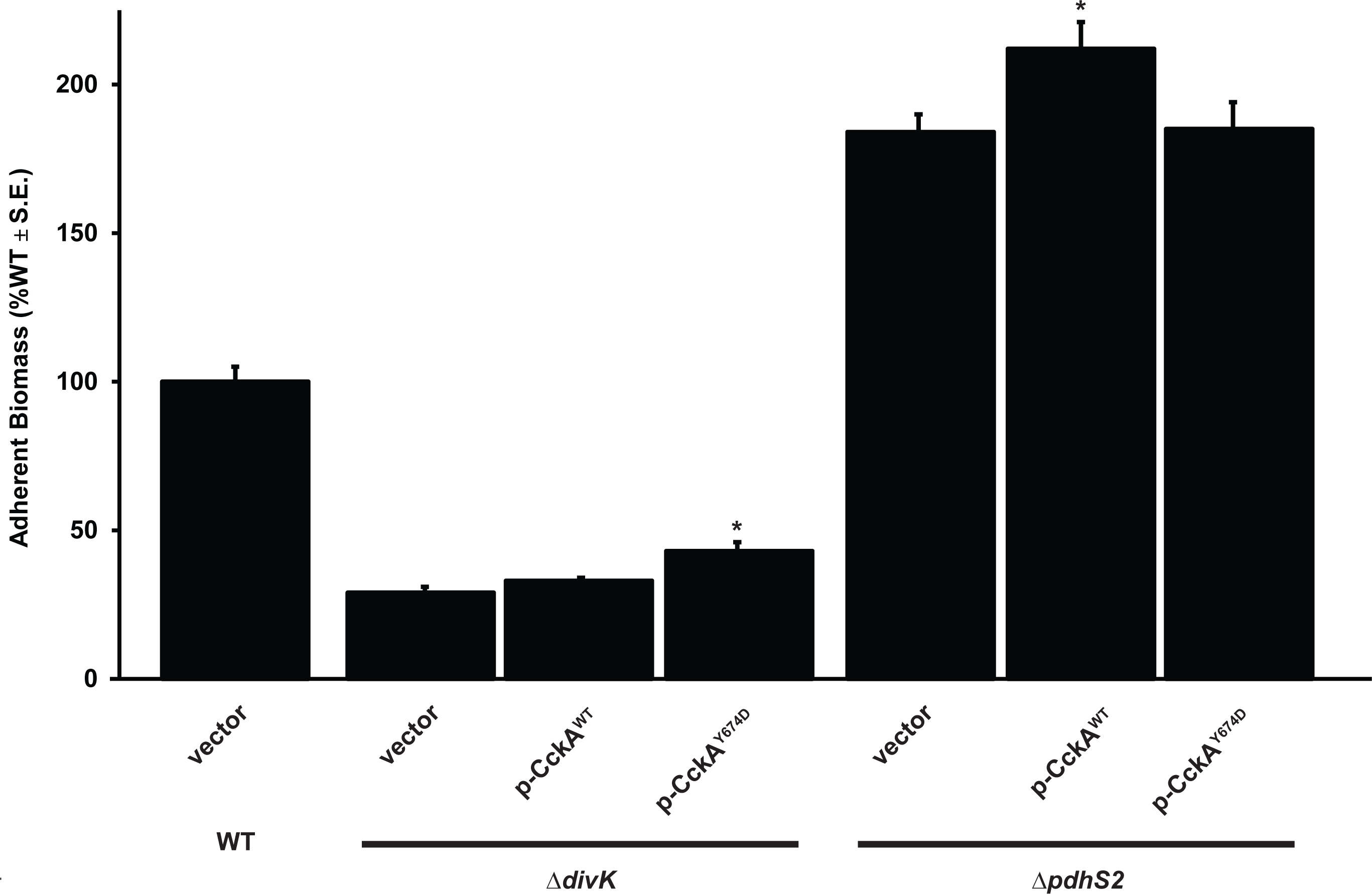
A kinase-locked allele of CckA fails to suppress the PdhS2-dependent biofilm phenotype. Biofilm formation was evaluated in the indicated strains as described in Figure 2. (^*^) = *P* < 0.05 compared to background strain carrying vector alone. Statistical significance was determined using Student’s *t* test.

**Figure S4.**
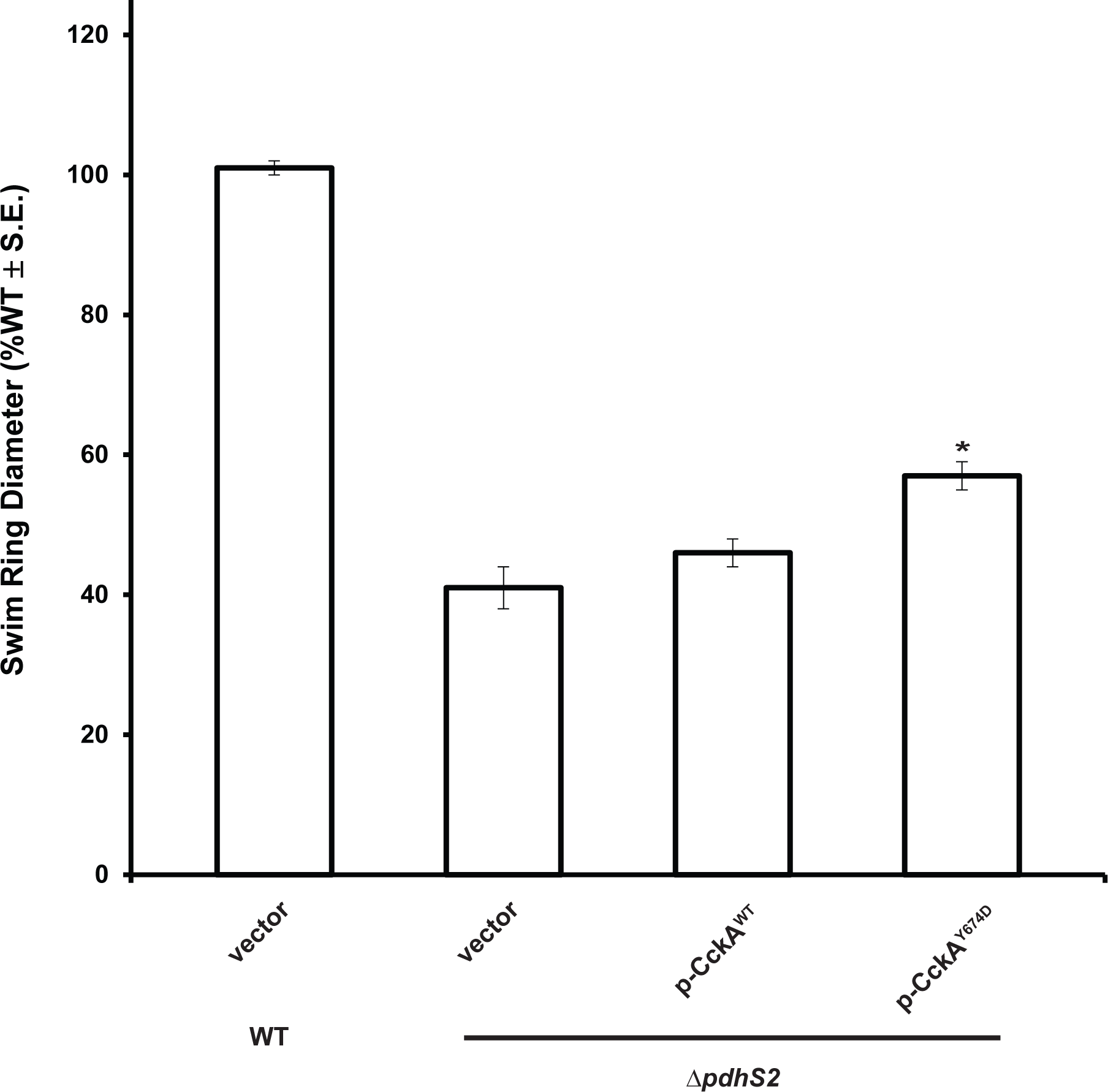
A kinase-locked allele of CckA fails to suppress the PdhS2-dependent swimming motility phenotype. Swimming motility was evaluated in the indicated strains as described in Figure 2. (^*^) = *P* < 0.001 compared to background strain carrying vector alone. Statistical significance was determined using Student’s *t* test.

**Figure S5.**
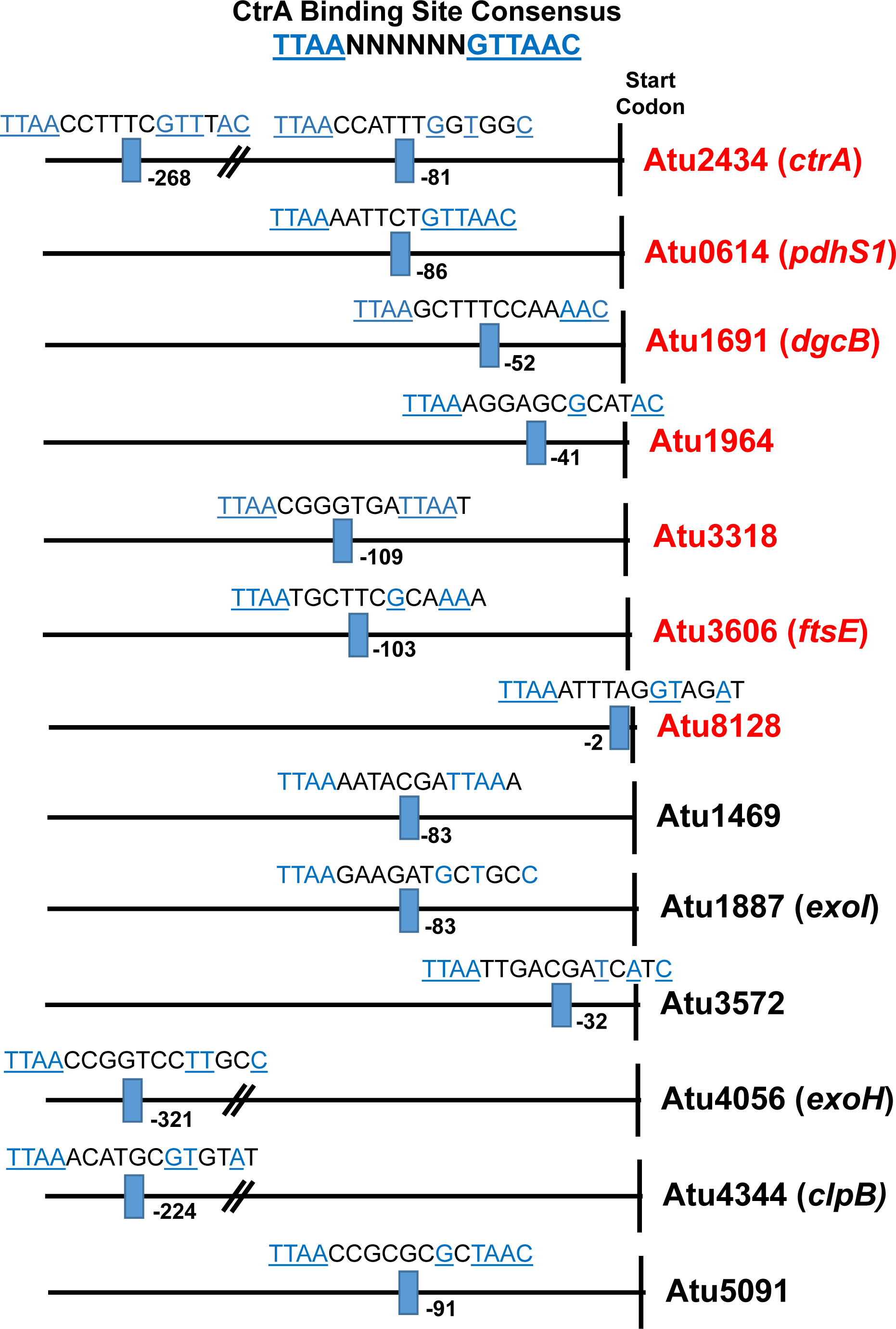
Predicted CtrA-dependent promoters bearing one or more CtrA binding motifs. Upstream regions from genes whose expression is increased (red) or decreased (black) in the Δ*pdhS2* mutant background relative to the wild-type background (microarray or *lacZ* fusion data). Predicted CtrA binding sites, as defined in the main text, are indicated. For genes in operons, the most upstream gene that has a predicted CtrA box is shown, even if this upstream gene did not make the expression cutoff for Table 2 (e.g. Atu4344 and Atu 8282). Only those genes with a predicted full CtrA binding site are shown.

**Figure S6.**
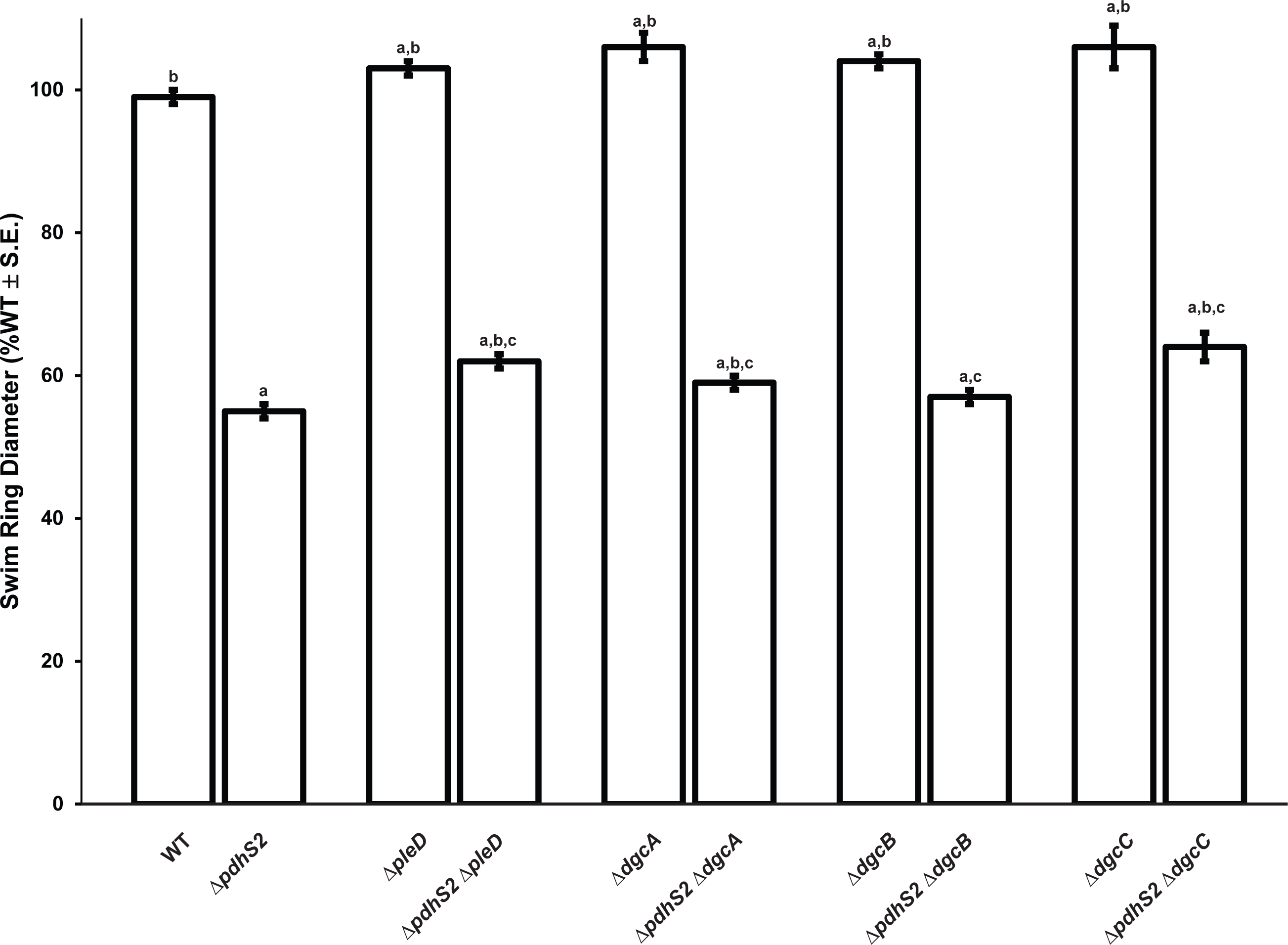
PdhS2 regulation of swimming motility is independent of diguanylate cyclase activity. Swimming motility of the wild-type (WT) and indicated mutant strains was evaluated as described in Figure 2. *P* < 0.05 compared to WT (^a^), Δ*pdhS2* (^b^), or corresponding diguanylate cyclase (^c^).

**Figure S7.**
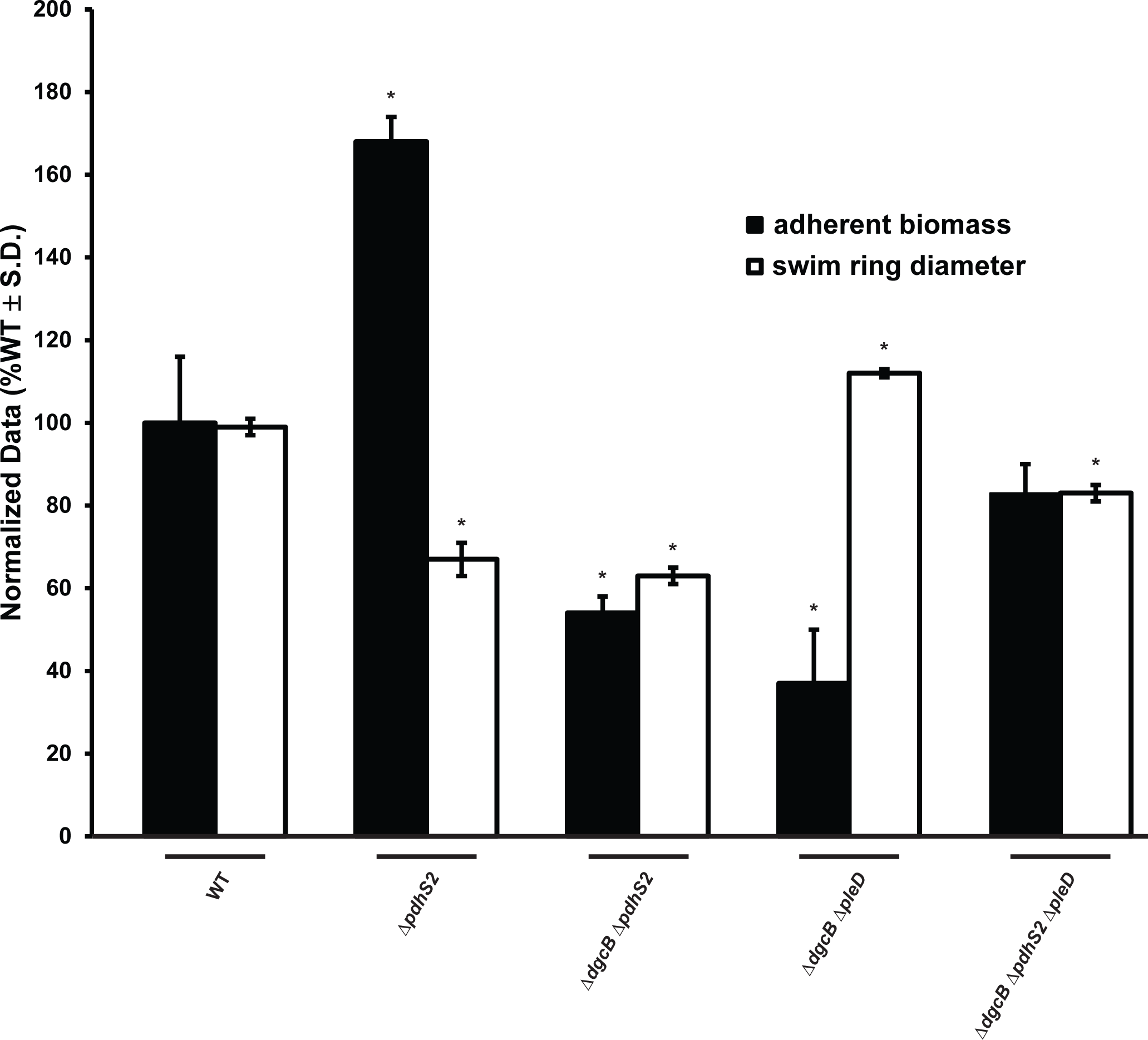
Loss of *pdhS2* enhances biofilm formation in the absence of both *dgcB* and *pleD*. Biofilm formation and swimming motility was evaluated in the wild-type (WT) and indicated mutant strains as described in Figure 2. (^*^) = *P* < 0.05 compared to the wild-type background. Statistical significance was determined using Student’s *t* test.

**Figure S8.**
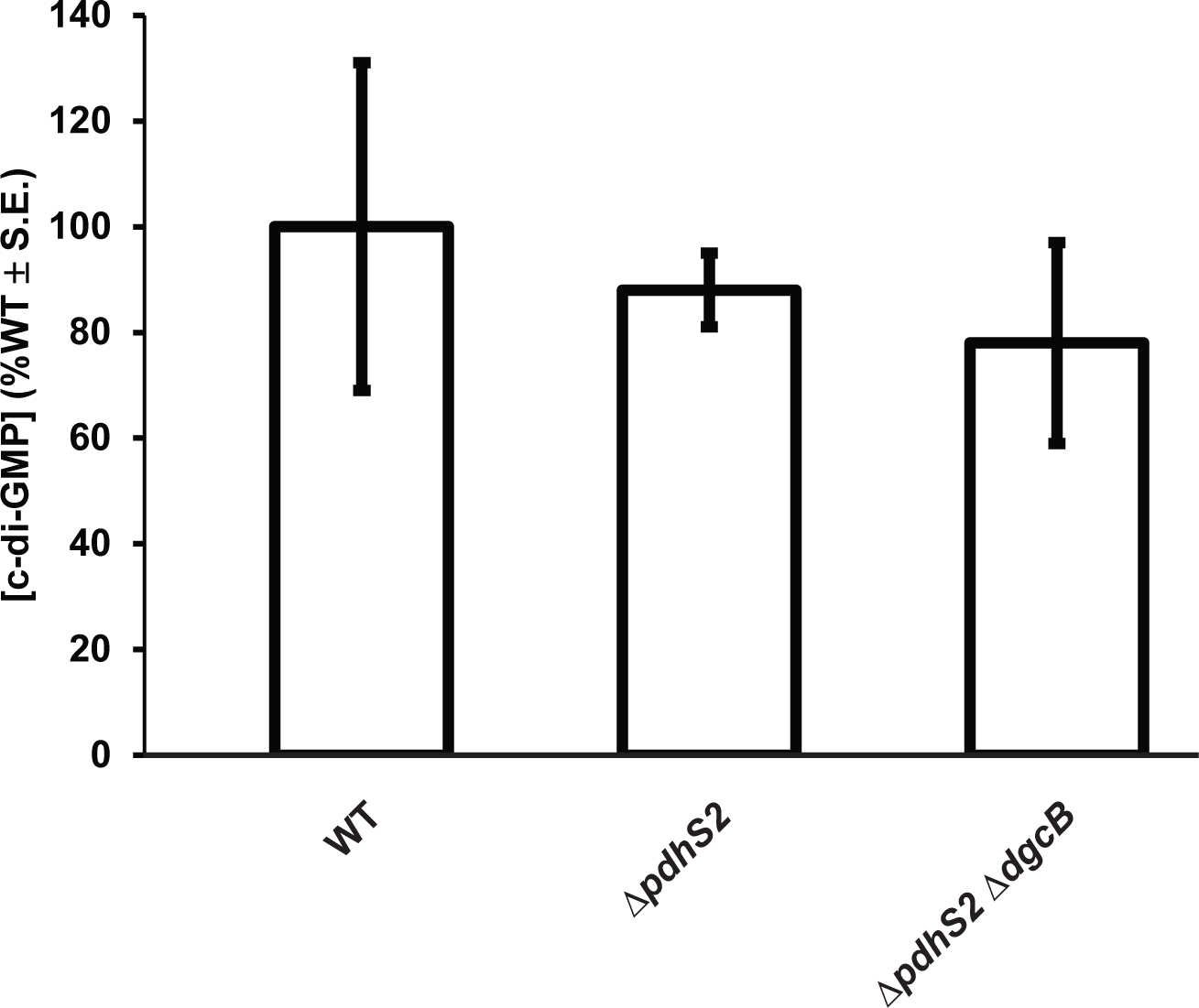
PdhS2 does not affect global levels of cyclic-di-GMP. Cyclic-di-GMP levels were measured in whole cell extracts from equivalent ODs of the indicated strains. Data are from three independent experiments (N = 3).

**Figure S9.**
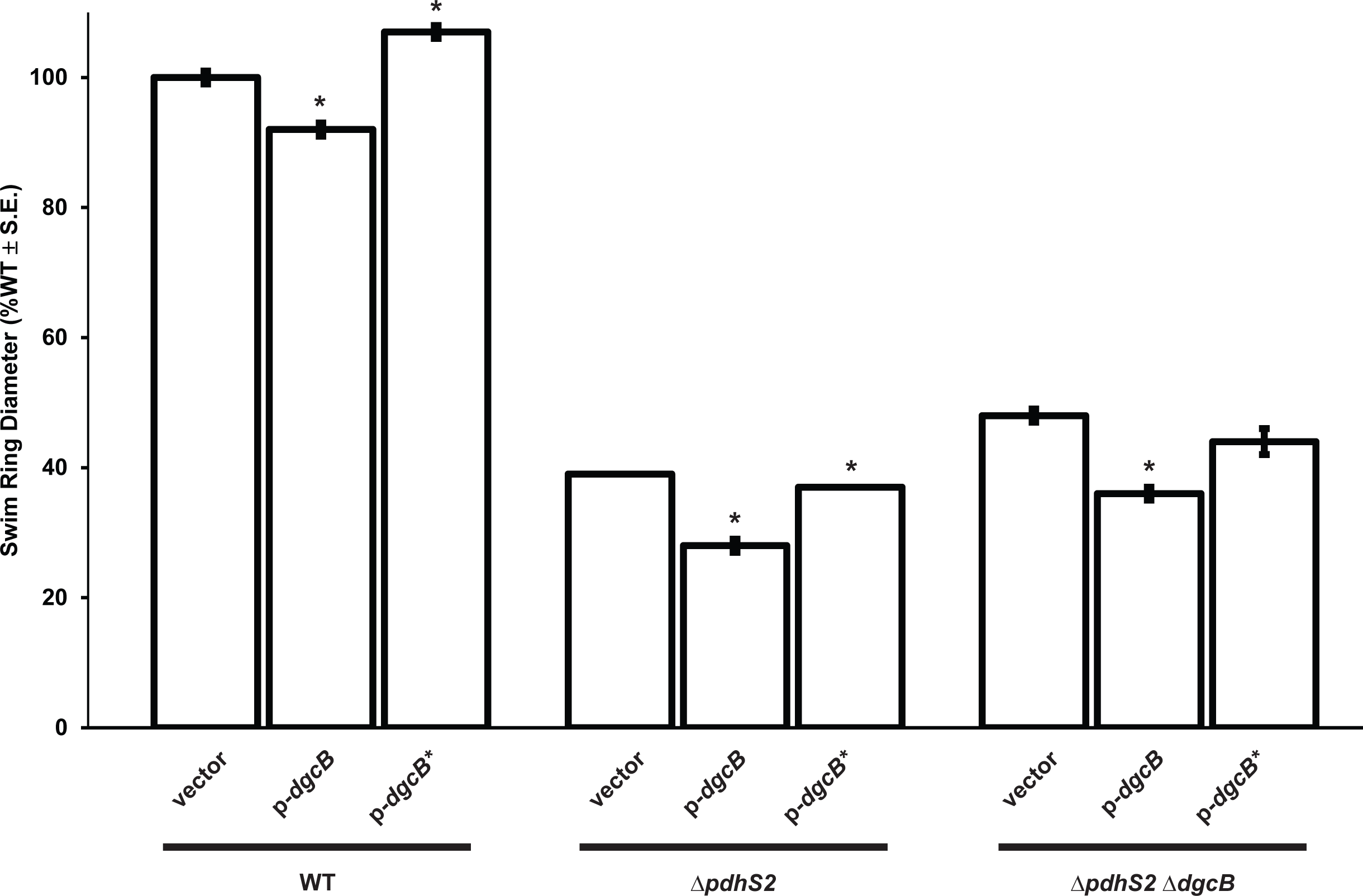
A catalytically inactive DgcB does not affect swimming motility. The effect on swimming motility of plasmid-borne expression of wild-type *dgcB* (p-*dgcB*) or a catalytic mutant allele of *dgcB* (p-*dgcB*\*) was evaluated. Expression of each *dgcB* allele was driven by the *P*_*lac*_ promoter. Biofilm formation was evaluated as described in Figure 2. (*) = *P* < 0.05 compared to vector alone.

**Figure S10.**
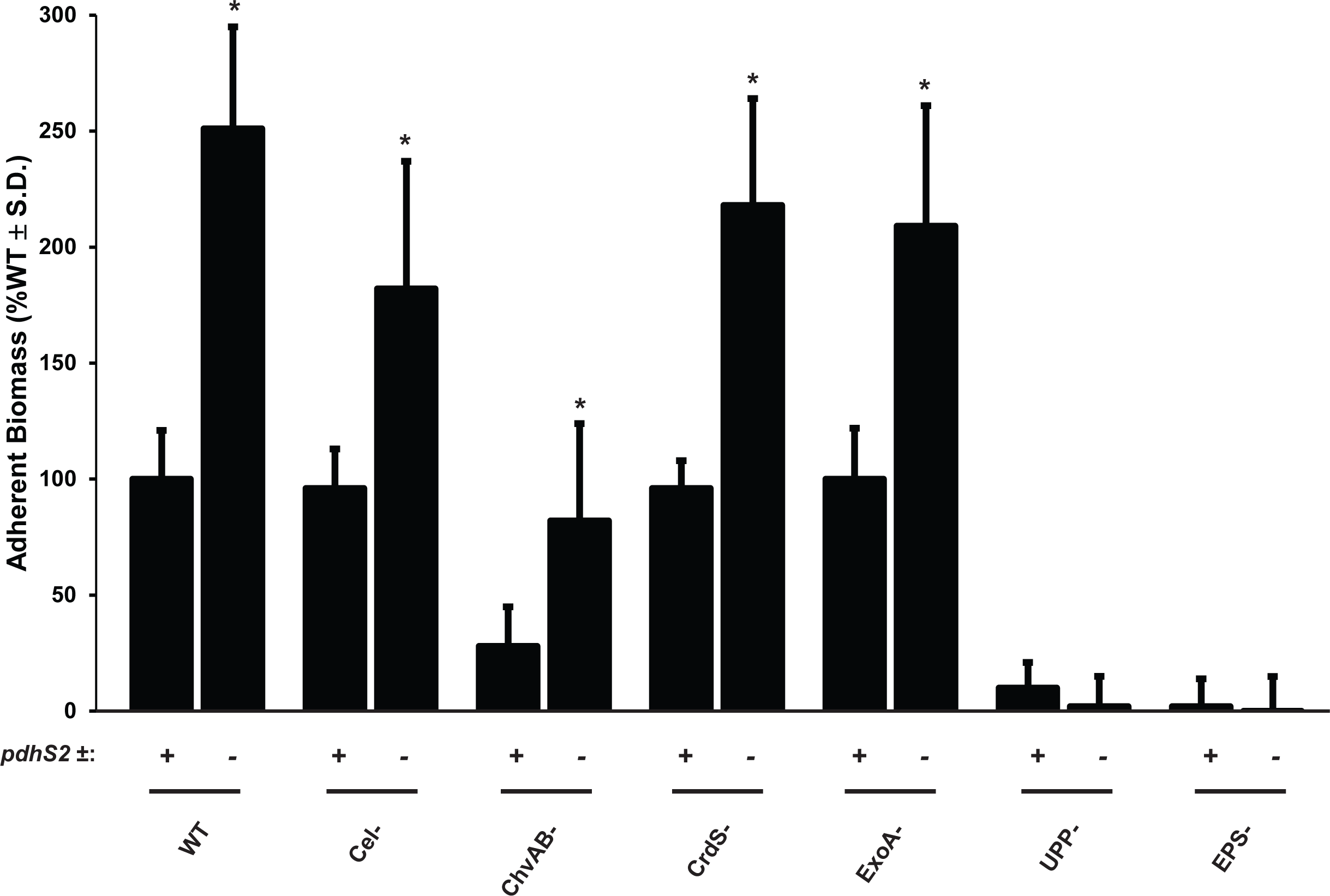
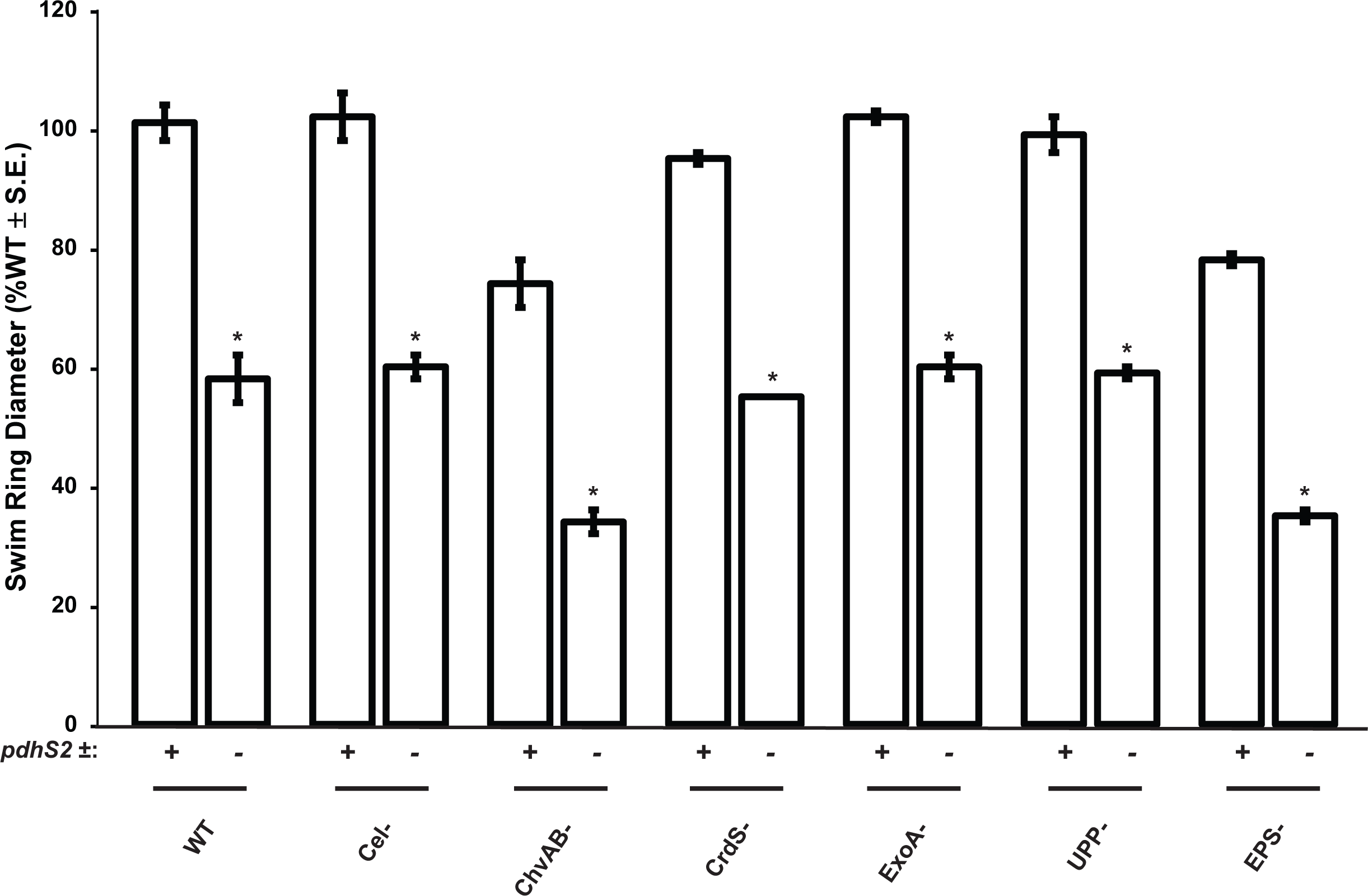
The unipolar polysaccharide is required for PdhS2-dependent biofilm formation. (A) Biofilm formation was evaluated in the presence (+) or absence (−) of *pdhS2* in combination with the indicated polysaccharides. WT = wild-type, Cel^−^ = cellulose mutant, ChvAB^−^ = cyclic-β-glucan mutant, CrdS^-^ = curdlan mutant, ExoA^−^ = succinoglycan mutant, UPP^−^ = unipolar polysaccharide mutant, EPS^−^ = mutant lacking all of the above polysaccharides. (B) Swimming motility was evaluated in the same strains as in (A). (*) = *P* < 0.05 compared to background strain. Statistical significance was determined using Student’s *t* test.

**Table S1.**
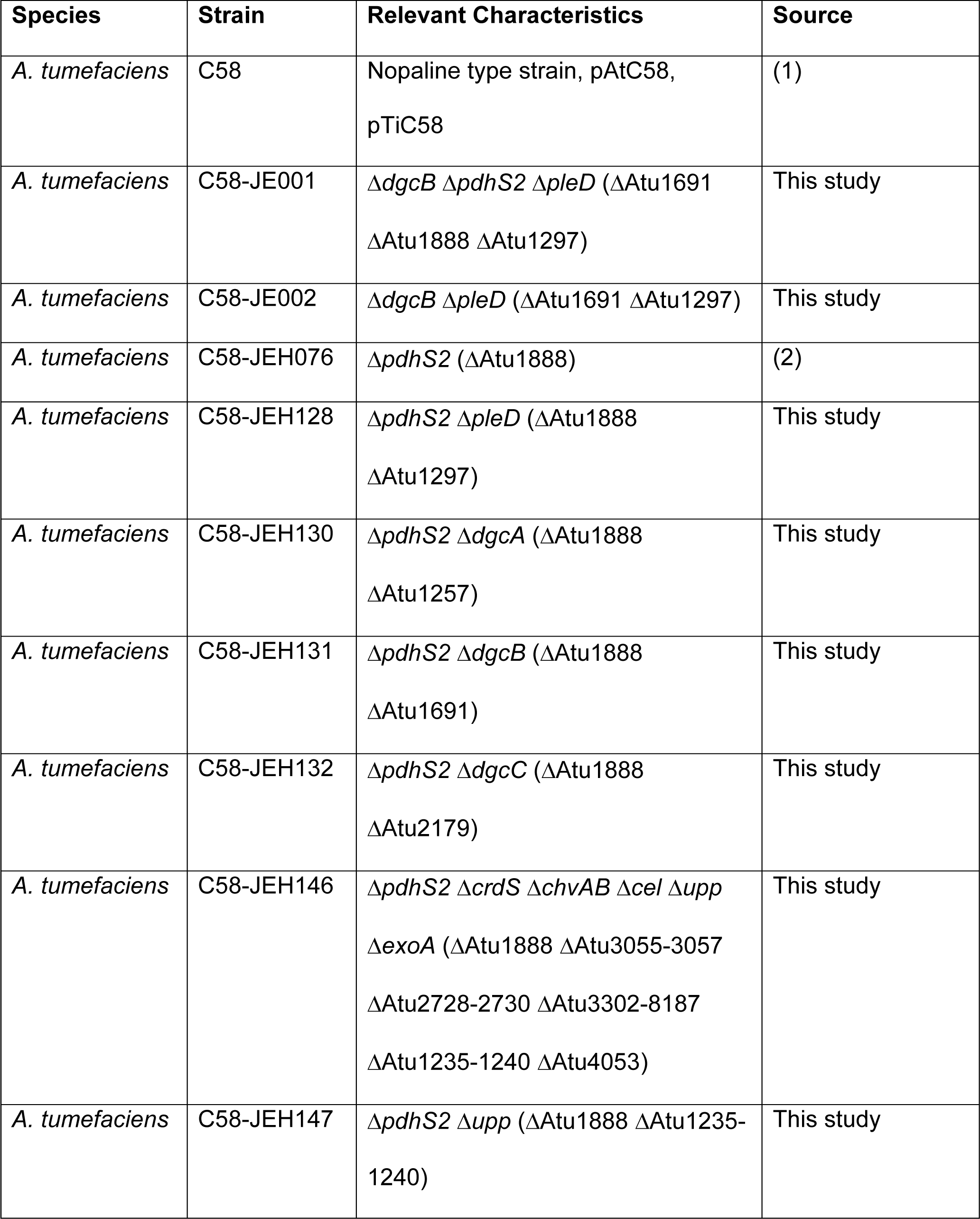
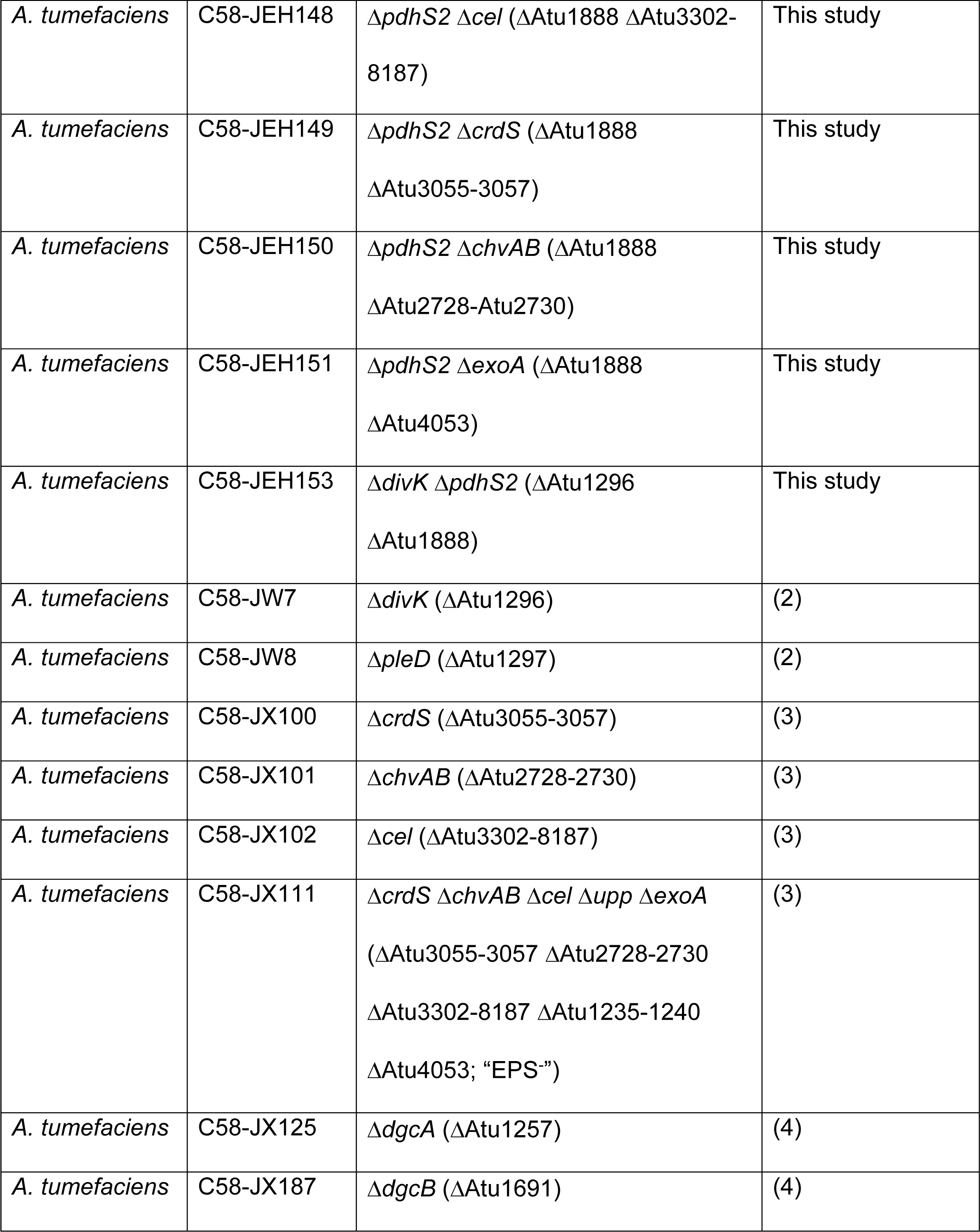
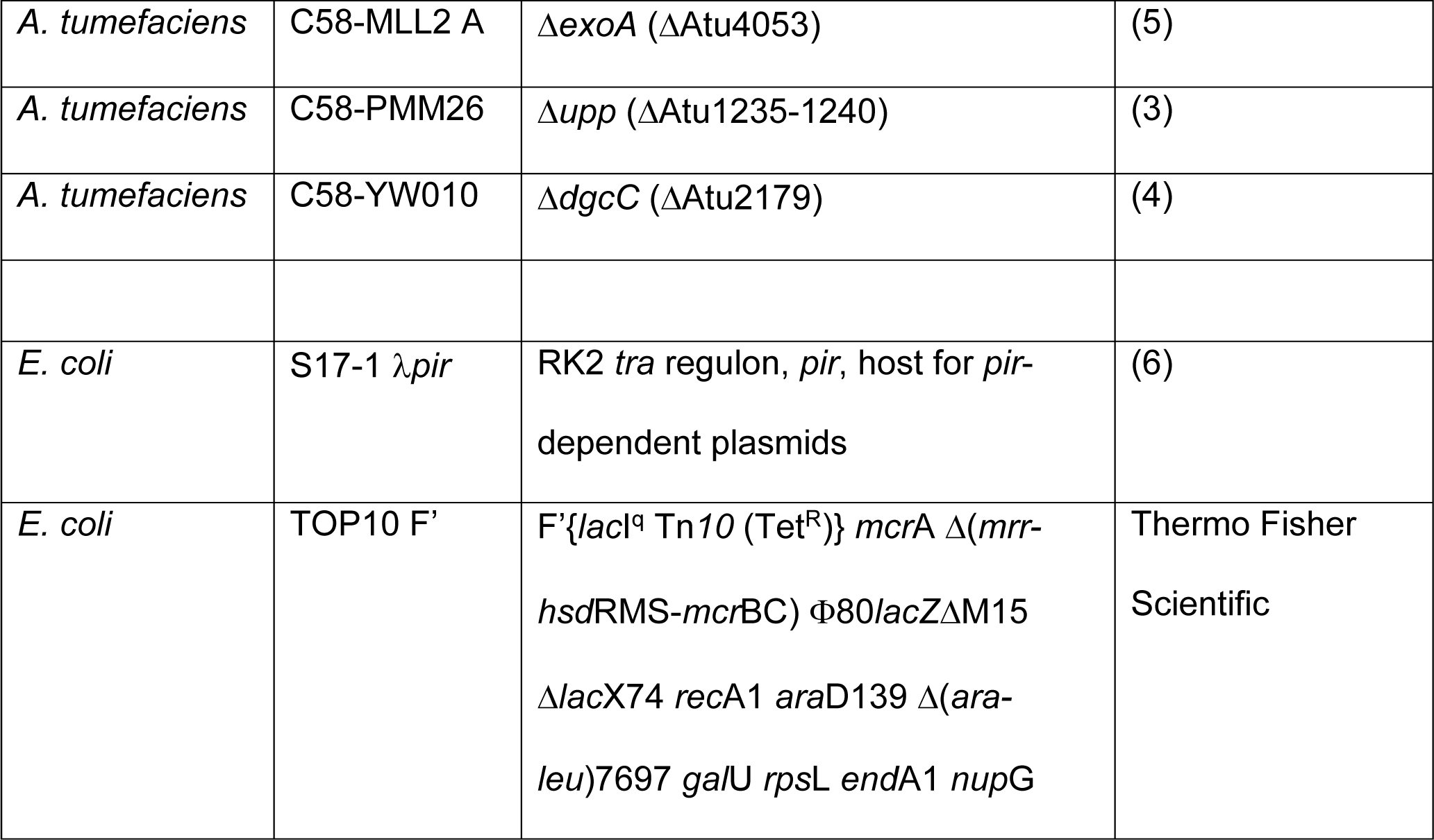
Strains used in this study

**Table S2.** Plasmids used in this study

| Plasmid name | Relevant characteristics | Source |
| --- | --- | --- |
| pGEM-T Easy | PCR cloning vector; Amp<sup>R</sup> | Promega |
| p*lacZ*/290 | Broad host range plasmid carrying promoterless *lacZ* for transcriptional fusions; Tet<sup>R</sup> | (7) |
| pNPTS138 | ColE1 origin; *sacB*; Km<sup>R</sup> | gift of M. Alley |
| pRA301 | Broad host range plasmid carrying promoterless *lacZ* for translational fusions; Spec<sup>R</sup> | (8) |
| pSRKGm | Broad host range vector containing $P_{lac}$; *lacI<sup>q</sup>*; *lacZ*$\alpha^+$; Gm<sup>R</sup> | (9) |
| pctrA290 | p*lacZ*/290 derivative with *C. crescentus ctrA* promoter | (10) |
| pDC001 | pGEM-T Easy with full-length *pdhS2*<sup>(CA811-812GC, A823G)</sup> (PdhS2<sup>His271A,Thr275Ala</sup> mutant) | This study |
| pDC002 | pSRKGm with full-length<br>*pdhS2*<sup>(CA811-812GC, A823G)</sup><br>(PdhS2<sup>His271A,Thr275Ala</sup><br>mutant) | This study |
| pGZ22 | *placZ*/290 derivative with<br>*C. crescentus ccrM*<br>promoter | (11) |
| pJEH010 | pSRKGm with full-length<br>wild-type *cckA* | (2) |
| pJEH021 | pGEM-T Easy with full-length *pdhS2* | (2) |
| pJEH026 | pSRKGm with full-length<br>*pdhS2* | (2) |
| pJEH030 | pSRKGm with full-length<br>Y674D *cckA* allele | (2) |
| pJEH040 | pNPTS138 derivative with<br>*pdhS2* SOE deletion<br>fragment | (2) |
| pJEH052 | pGEM-T Easy with *pdhS2*<br>lacking a stop codon | This study |
| pJEH053 | pGEM-T Easy with<br>*gfpmut3* | This study |
| pJEH054 | pGEM-T Easy with *divJ*<br>lacking a stop codon | This study |
| pJEH060 | pSRKGm with a<br>*pdhS2::gfpmut3*<br>translational fusion | This study |
| pJEH078 | pSRKGm with a<br>*divJ::gfpmut3* translational<br>fusion | This study |
| pJEH091 | pGEM-T Easy with full-length *pdhS2*<sup>(CA811-812GC)</sup><br>(PdhS2<sup>His271Ala</sup> mutant) | This study |
| pJEH092 | pSRKGm with full-length<br>*pdhS2*<sup>(CA811-812GC)</sup><br>(PdhS2<sup>His271Ala</sup> mutant) | This study |
| pJEH099 | pGEM-T Easy with full-length *pdhS2*<sup>(A823G)</sup><br>(PdhS2<sup>Thr275Ala</sup> mutant) | This study |
| pJEH102 | pSRKGm with full-length<br>*pdhS2*<sup>(A823G)</sup><br>(PdhS2<sup>Thr275Ala</sup> mutant) | This study |
| pJEH113 | pGEM-T Easy with A.<br>*tumefaciens ccrM*<br>promoter | This study |
| pJEH115 | pGEM-T Easy with A.<br>*tumefaciens ctrA* promoter | This study |
| pJEH119 | pGEM-T Easy with A.<br>*tumefaciens pdhS1*<br>promoter | This study |
| pJEH121 | pRA301 with A.<br>*tumefaciens ccrM*<br>promoter | This study |
| pJEH122 | pRA301 with A.<br>*tumefaciens ctrA* promoter | This study |
| pJEH124 | pRA301 with A.<br>*tumefaciens pdhS1*<br>promoter | This study |
| pJEH133 | pRA301 with mutated A.<br>*tumefaciens dgcB*<br>promoter | This study |
| pJS70 | p*lacZ*/290 derivative with<br>*C. crescentus pilA*<br>promoter | (12) |
| pJX158 | pRA301 with A.<br>*tumefaciens* Atu3318<br>promoter | (4) |
| pJX162 | pRA301 with *A. tumefaciens dgcB* promoter | (4) |
| pJX520 | pSRKGm with full-length *dgcB* | (4) |
| pJX521 | pSRKGm with full-length *dgcB*<sup>A767C, A770C</sup> (DgcB<sup>EE256-257AA</sup> mutant) | (4) |
| pJX802 | pNPTS138 derivative with *dgcB* SOE deletion fragment | (4) |
| pJZ383 | pPZP201 derivative with *P<sub>tac</sub>::gfpmut3*; Spec<sup>R</sup> | (13) |

**Table S3.**
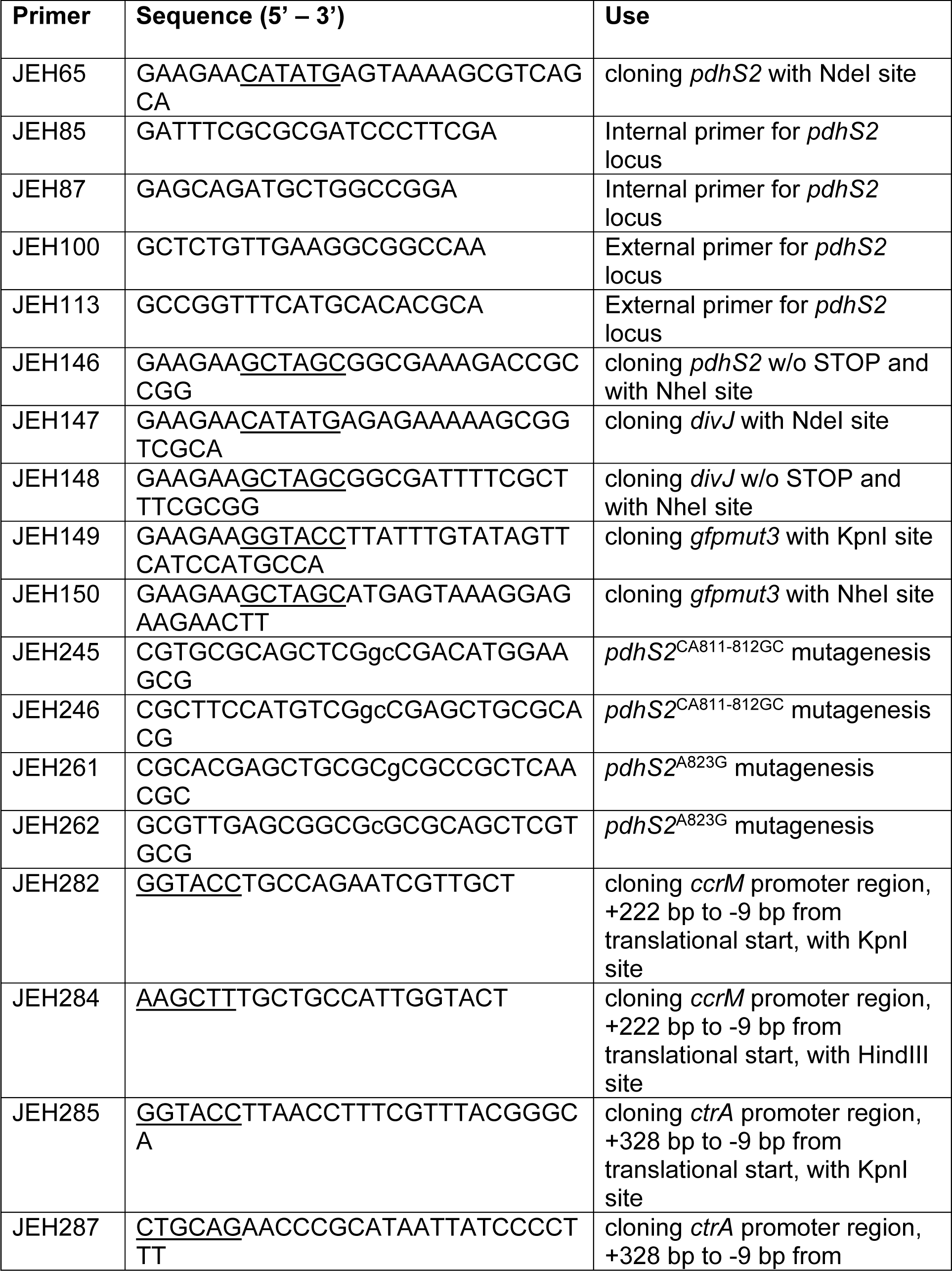
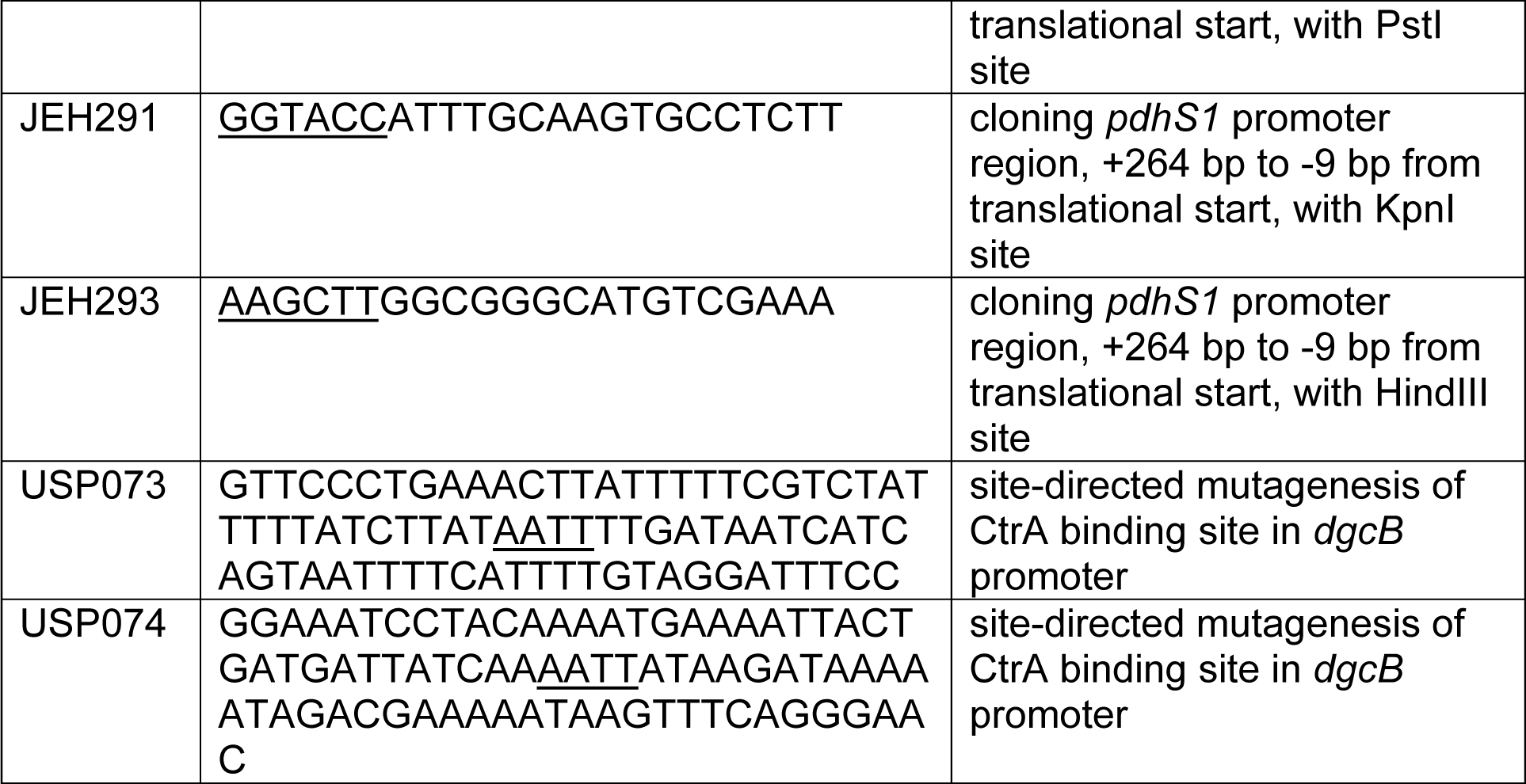
Primers used in this study

